# Cracking the black box of deep sequence-based protein-protein interaction prediction

**DOI:** 10.1101/2023.01.18.524543

**Authors:** Judith Bernett, David B. Blumenthal, Markus List

## Abstract

Identifying protein-protein interactions (PPIs) is crucial for deciphering biological pathways. Numerous prediction methods have been developed as cheap alternatives to biological experiments, reporting surprisingly high accuracy estimates. We systematically investigated how much reproducible deep learning models depend on data leakage, sequence similarities, and node degree information, and compared them to basic machine learning models. We found that overlaps between training and test sets resulting from random splitting lead to strongly overestimated performances. In this setting, models learn solely from sequence similarities and node degrees. When data leakage is avoided by minimizing sequence similarities between training and test set, performances become random. Moreover, baseline models directly leveraging sequence similarity and network topology show good performances at a fraction of the computational cost. Thus, we advocate that any improvements should be reported relative to baseline methods in the future. Our findings suggest that predicting protein-protein interactions remains an unsolved task for proteins showing little sequence similarity to previously studied proteins, highlighting that further experimental research into the “dark” protein interactome and better computational methods are needed.

## Introduction

Proteins carry out essential biological functions, many of which require proteins to act jointly or to form complexes. Hence, identifying all pairwise interactions of proteins is an essential systems biology challenge toward understanding biological pathways and their dysregulation in diseases. Several technologies (e.g., yeast-2-hybrid screens, affinity purification mass-spectrometry) have been developed to unravel individual PPIs, yielding large-scale PPI networks^1^. As it is not feasible to study all protein pairs exhaustively, a plethora of computational methods have been developed to predict PPIs as a binary classification task. Such methods often use only sequence information in various machine learning (ML) strategies, ranging from classical support vector machines to the most complex deep learning (DL) architectures currently conceivable^2–21^. These DL methods typically report phenomenal prediction accuracies in the range of 95%-99%.

Though it was criticized that only a few of these methods have source code available and are reproducible^2,22^, it has not yet been examined systematically whether and how such results are possible. Since proteins interact in 3D space, predicting an interaction should implicitly consider the 3D structure of complexes, binding pockets, domains, surface residues, and binding affinities. However, predicting protein 3D structure from sequence is an infamously hard problem area in which only recently AlphaFold2 had made a tremendous leap using a very complex model architecture and vast resources^23^. Moreover, predicting the structure of multi-chain complexes observed in PPIs remains an open challenge^24^. In light of this, the observed high accuracies for predicting PPIs from sequence information alone seem dubious.

Few studies shed light on the phenomenal accuracies reported for deep sequenced-based PPI prediction approaches: Almost all PPI datasets used for evaluating such approaches are randomly split into train and test sets using cross-validation. Park & Marcotte^25^ showed that this causes an inflation of prediction performance due to training data leakage^25–27^. Upon random splitting, the same proteins occur both in the train and the test set, such that these sets are no longer independent^28^. For an extensive definition of data leakage, see^29,30^. To quantify the effect of data leakage, Park & Marcotte^25^ proposed three classes C1 (both proteins in a test pair occur in training), C2 (only one protein in a test pair occurs in training), and C3 (no overlap between training and test). Prediction accuracies usually drop significantly between C1 and C2 as well as between C2 and C3. It has also been shown that when datasets contain sequences with high pairwise sequence similarities, models overfit and accuracies are overestimated, giving a wrong impression of the state of the field^22,26,27^. Hamp & Rost^26^, therefore, extended the requirements by demanding that, for C3, no test protein should be sequence-similar to a training protein (for C2 only one, for C1 both), and obtained similar results as Park & Marcotte. Furthermore, Chatterjee al.^31^ have recently shown that deep learning methods for protein-ligand prediction use degree information as shortcuts instead of learning from sequence features. A baseline model using only topology information performs equally well for that task. They reveal that protein hubs have disproportionally more positive annotations and that most proteins and ligands either have almost no positive or almost no negative annotations, which the methods leverage. In addition to positive examples, it is crucial to add sufficiently realistic negative examples to both train and test sets. These can be randomly sampled by choosing protein pairs not reported as PPIs in public databases. To avoid using false negatives, it is common practice to choose proteins that are not annotated to the same cellular compartment and are thus not expected to interact in a cell. However, Ben-Hur & Noble^32^ have shown that this approach makes the learning problem considerably easier than it is in reality. Moreover, protein pairs are tested for interaction in an artificial system^33^ and databases thus frequently include interactions of pairs annotated to different cellular locations.

In this work, we systematically examine three scenarios that might explain why sequence-based models correctly predict whether two proteins *p*_1_ and *p*_2_ interact:

- Explanation 1: The models can detect patterns in the sequences of *p*_1_ and *p*_2_ that are responsible for whether they can interact (e.g., matching binding sites, domains, motifs).
- Explanation 2: The models utilize node degree information shortcuts that individually explain whether the protein interacts. Based on these individual tendencies, they predict the interactions (e.g., if, in the training fold, *p*_1_ only appears in interactions and never in the negative set, protein pairs from the training fold that involve *p*_1_ will likely be predicted as interacting).
- Explanation 3: The models merely check whether *p*_1_ and *p*_2_ are similar to protein sequences 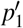and 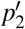from the training set and make a prediction based on the interactions of 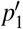 and 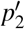(e.g., if 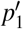interacts with 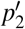, *p*_1_ probably interacts with *p*_2_ as well).

While, for many methods, the authors (implicitly) assume that their method’s excellent prediction performance can be attributed to Explanation 1, we hypothesize that Explanation 2 and Explanation 3 are the actual drivers of high prediction accuracy. To investigate Explanation 2, we randomized the positive input network in the training data (but not in the test data) via degree-preserving rewiring. Hence, each protein’s node degree is preserved, but the edges are no longer biologically plausible. If node degree information shortcuts indeed drove prediction performance, we would expect only a moderate drop in the test accuracy. Additionally, we incorporated two baseline methods (harmonic function^34^ and local and global consistency algorithm^35^) that exclusively utilize network topology to infer if two proteins interact. To investigate Explanation 3, we carried out a two-fold strategy: On the one hand, we compared deep sequenced-based PPI prediction approaches against SPRINT^36^ — an algorithmic PPI prediction approach based on local sequence alignment — as well as against simple baseline ML models which, by design, have access only to sequence similarity information. If sequence similarity indeed were a main driver of prediction performance, we would expect our baselines to achieve similar performances as the state-of-the-art DL methods. On the other hand, we partitioned the proteome into two blocks such that inter-block sequence similarities are minimized and selected PPIs for train and test sets from different blocks of the partition. This ensures that sequence similarity patterns learned during training cannot be leveraged at test time. If Explanation 3 were valid, we would expect a significant drop in prediction performance.

We conducted our analyses for six deep sequence-based PPI prediction methods^2,4,13,20,21^, which we trained and tested on the same seven publicly available and commonly used datasets (two datasets with yeast PPIs^12,37^ and five with human PPIs^2,11,20,38^). Our results show that training data leakage can fully explain the excellent accuracies reported in the literature. More specifically, if pairwise sequence similarities are minimized between disjoint training and test sets, performance is random, proving that sequence similarity and node degree are the only relevant features in current sequence-based PPI prediction methods. Finally, we generated a gold standard dataset to enable data-leakage-free validation of future PPI prediction methods.

## Results

### Overview

We reviewed the literature for PPI prediction methods and their underlying datasets (Supplemental Table S1). For most of the 32 methods we found, extraordinary prediction performances are reported. However, source code is available only for twelve of them, emphasizing the reproducibility crisis in ML-based science^29^. Since we focused on understanding how sequence information contributes to DL-based PPI prediction, we selected methods that we managed to reproduce with reasonable effort and which rely exclusively on sequence information. This reduced the number of DL methods to Richoux-FC, Richoux-LSTM^2^, DeepFE^13^, PIPR^4^, D-SCRIPT^20^, and Topsy-Turvy^21^.

For testing how much can be predicted from topology alone, we incorporated two node classification algorithms (harmonic function^34^, local and global consistency^35^), which operate on the line graphs of the input networks. Additionally, we tested SPRINT^36^, a fast algorithmic method that uses only sequence similarities of protein pairs to predict PPIs. We also included two baseline ML models (Random Forest, SVM) that used dimensionality-reduced (PCA, MDS, node2vec) sequence similarity vectors as input for each protein. These baseline methods allowed us to assess the benefit of DL and to test the hypothesis that sequence similarity alone is already sufficient to achieve good prediction performance. Pairwise sequence similarities were pre-computed by SIMAP2^39^. Although sequence similarities were the only input feature, we note that the methods could learn node degrees implicitly during training. Supplemental Figure S1 depicts a schematic overview of the methods’ principles.

We tested the methods on popular yeast and human datasets, the dataset used to validate the D-SCRIPT method (D-SCRIPT UNBALANCED)^20^, and the two datasets by Richoux et al.^2^ (see Table 1 for an overview). The latter two datasets were included because of their size and the unique generation of the strict test dataset, which was designed to be free from hub biases. D-SCRIPT UNBALANCED was included because it is deliberately unbalanced (one to ten positive versus negative annotations) to better reflect the underlying label distribution. The two Richoux datasets were created from a larger dataset consisting of PPIs annotated in Uniprot, which we later used for the partitioning task and refer to as RICHOUX-UNIPROT. All datasets were cleaned from duplicates and balanced except for D-SCRIPT UNBALANCED). Because of GPU restrictions, we created length-restricted datasets for D-SCRIPT and Topsy-Turvy, in which each protein had between 50 and 1000 amino acids. The original datasets were split 80/20 into training/test except for RICHOUX-REGULAR, RICHOUX-STRICT, and D-SCRIPT UNBALANCED, which were already split into training/(validation/) test. Since we only used default hyperparameters, we did not need a validation set and added the validation to training for these two datasets. We chose the random 80/20 split since most reviewed methods report the mean accuracy of five-fold cross-validation^4–8,11,18,38,41,42^ or a random hold-out test set^2,3,12,13,37,43–46^. As many datasets are rather small, those models which were developed for larger datasets (e.g., D-SCRIPT and Topsy-Turvy) could be prone to overfitting. We therefore also tested how early stopping influences the results of all deep learning methods. For picking the best model from the epochs, we randomly took 10% of the training set as validation set in this setting.

**Table 1:** Overview of datasets. *n* denotes the overall number of samples in the datasets (after balancing and removal of duplicates), i.e., the number of PPIs plus the number of randomly sampled non-edges. *n*_restr._ is size of the length-restricted datasets where both proteins of each (non-)interaction have between 50 and 1000 amino acids. These datasets were used for D-SCRIPT and Topsy-Turvy. *n*_method_ is number of methods found in our literature review that were tested on the respective datasets. MRA denotes the median reported accuracy of the *n*_method_ methods tested on the respective datasets. D-SCRIPT and Topsy-Turvy only reported auPR and AUC on their dataset.

| Dataset | Organism | $n$ | $n_{\text{restr.}}$ | $n_{\text{method}}$ | MRA (in %) |
| --- | --- | --- | --- | --- | --- |
| HUANG<sup>40</sup> | Human | 6690 | 4758 | 4 | 98.43 |
| GUO<sup>37</sup> | Yeast | 11 162 | 8760 | 14 | 94.75 |
| DU<sup>12</sup> | Yeast | 34 512 | 27 356 | 5 | 92.50 |
| PAN<sup>38</sup> | Human | 62 962 | 44 920 | 5 | 96.82 |
| RICHOUX-REGULAR<sup>2</sup> | Human | 79 868 | 67 724 | 2 | 88.10 |
| RICHOUX-STRICT<sup>2</sup> | Human | 68 664 | 78 776 | 2 | 77.29 |
| D-SCRIPT UNBALANCED | Human | 426 492 | 426 283 | 2 | 0.5605 (auPR) |

Figure 1 provides an overview of our analyses. We first consider a random split into train and test set, which we expect to introduce data leakage (see Methods for details). To test how much the models learn from node degree only (Explanation 2), we next rewired the positive PPIs (edges in PPI networks) in all training folds. For this, we randomly re-assigned all edges but preserved the expected node degree for each protein, rendering the new positive PPI networks biologically meaningless (see Figure 6). Finally, we used the KaHIP^47^ method with length-normalized, pre-computed SIMAP2^39^ bitscores as input to partition the human and yeast proteomes into two blocks, *P*_0_ and *P*_1_, such that pairs of protein sequences from different blocks are dissimilar. Then, for each dataset, all PPIs (*p*_1_, *p*_2_) were assigned to three blocks *INTRA*_0_, *INTRA*_1_, and *INTER*, depending on whether *p*_1_ and *p*_2_ are both contained in *P*_0_, whether they are both contained in *P*_1_, or whether they are contained in different blocks of the partition {*P*_0_, *P*_1_}. If Explanations 2 and 3 apply, we expect a significant drop in accuracy if we train on the PPIs contained in *INTRA*_0_ and test on the ones contained in *INTRA*_1_, since there is no direct data leakage (*P*_0_∩*P*_1_ = 0/) and a minimized amount of indirect data leakage due to sequence similarity. If we train on *INTER* and test on *INTRA*_0_ or *INTRA*_1_, we expect a smaller drop in accuracy since there is data leakage.

**Figure 1:**
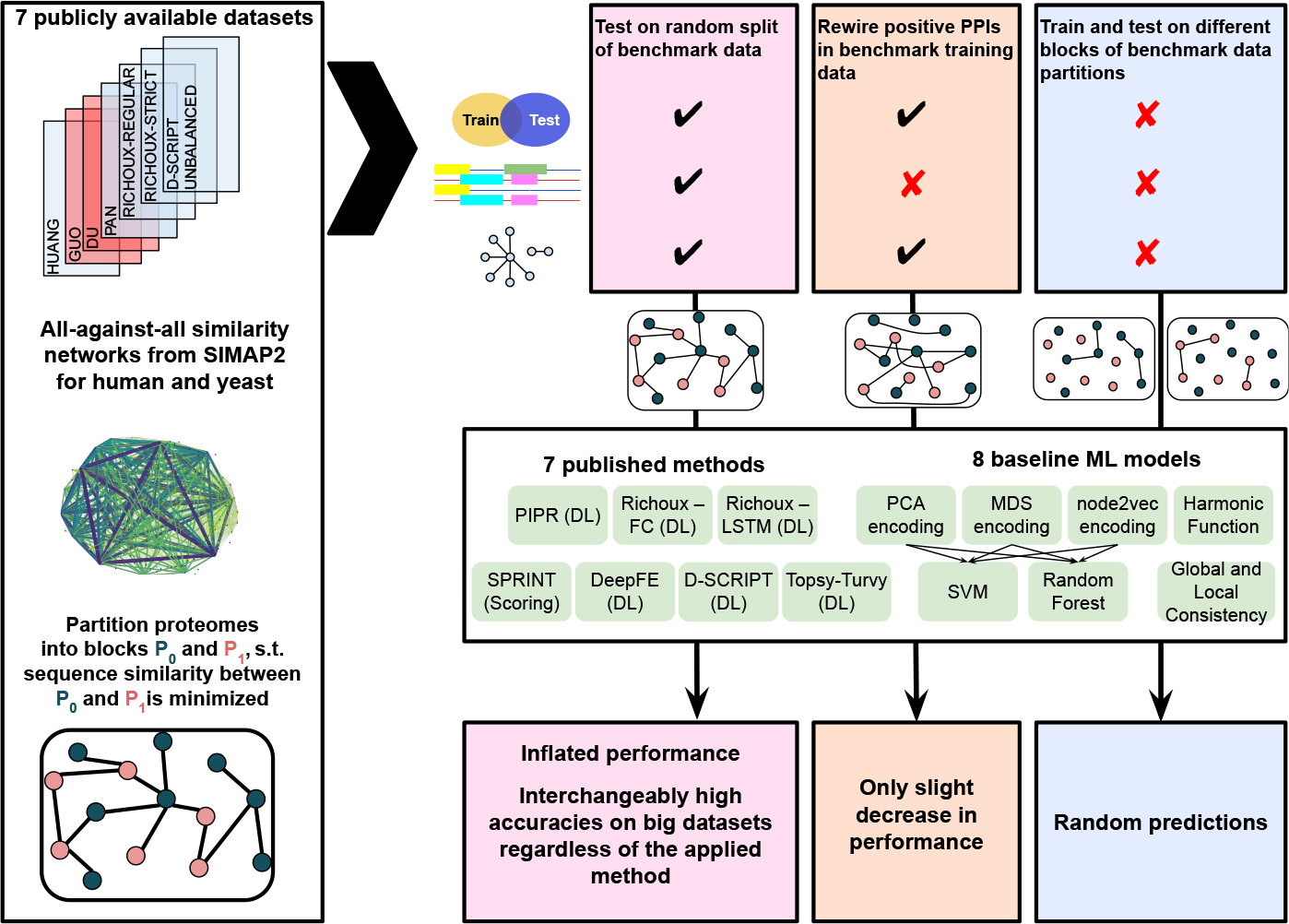
Overview of analyses. Seven publicly available datasets were used for testing seven published methods and eight basic ML models that use sequence similarities or topology as only input. We performed three tests to explain the phenomenal accuracies reported for DL methods: (1) We split the original data randomly into 80% train / 20% test, introducing data leakage through overlap of train and test proteins. Methods could learn from node degree biases and sequence similarities. This yielded inflated performance estimates and interchangeably high accuracies for DL and basic models on large enough datasets. (2) We rewired the positive train PPIs such that models could only learn from node degrees. Nevertheless, performance estimates only decreased slightly. (3) We partitioned the human and yeast proteomes into two blocks *P*_0_ and *P*_1_ such that proteins from different blocks have pairwise dissimilar sequences and assigned PPIs (*p*_1_, *p*_2_) to blocks *INTRA*_0_, *INTRA*_1_, and *INTER*, depending on whether *p*_1_ and *p*_2_ are both contained in *P*_0_ or *P*_1_, or fall into different blocks of the partition. When trained on *INTRA*_0_ and tested on *INTRA*_1_ (no overlap between train and test data, models could neither learn from sequence similarity nor from node degrees), all tested models predicted PPIs randomly.

## Results on randomly split original benchmark datasets

Figure 2 shows our results upon randomly splitting the original benchmark datasets into 80% train / 20% test. Since all published methods except for D-SCRIPT and Topsy-Turvy were reported to show close to perfect performances, we expected roughly comparable results across methods within each dataset. As larger data sets are more prone to data leakage, we further expected accuracy to increase with dataset size for random splitting.

**Figure 2:**
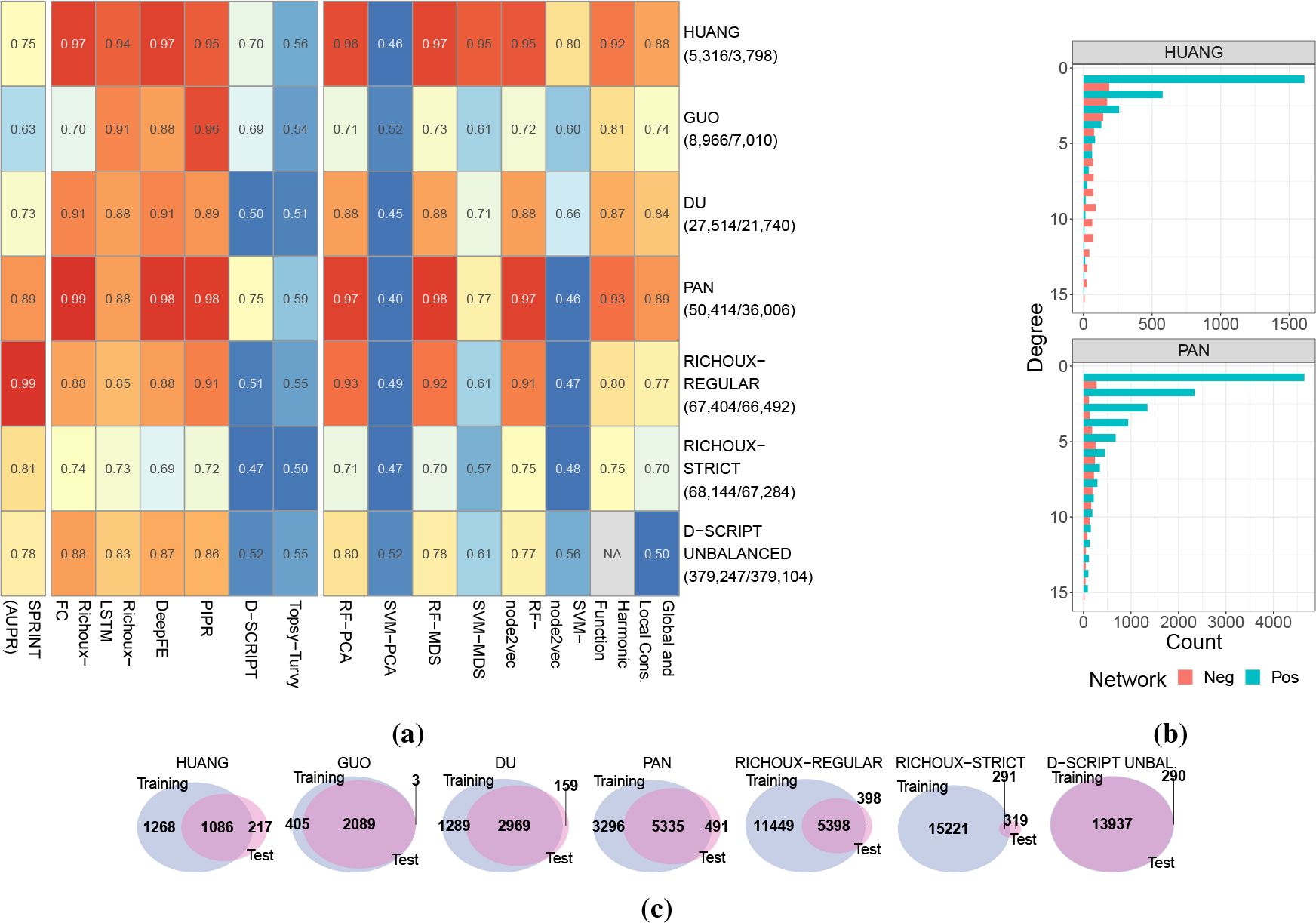
Results obtained on randomly split original benchmark datasets. **(a)** AUPRs on test sets for SPRINT, balanced accuracies for all other methods. Y-axis labels correspond to the dataset, the numbers of samples in the training data (all/restricted length) are shown in parantheses. AUPR values of SPRINT rise with the number of proteins in the dataset. Performances are exceptionally high on the HUANG and PAN dataset. For five out of eight baseline ML models, performances are comparable to the performances of the DL models. All performances drop significantly on RICHOUX-STRICT. **(b)** Node degree distributions of the positive and negative PPI datasets for HUANG and PAN. Only the node degrees of the positive PPIs follow a power law distribution. **(c)** Overlap of proteins occurring in the original training and test sets.

Comparing the results for all datasets except for the RICHOUX-STRICT dataset, we can see that the area under the precision-recall curve (AUPR) values of SPRINT rise with the number of unique proteins in the dataset (Supplemental Table S2). SPRINT’s AUPRs match the balanced accuracies of the DL models on large datasets, which shows that finding similar subsequences to predict PPIs is already sufficient to reach excellent performance measures when proteins between the train and test set are shared.

For almost all methods, performances are exceptionally high on the HUANG and PAN datasets. This phenomenon can be explained by looking at the node degree distributions of the positive and negative datasets (Figure 2b). While both distributions follow the power law for the other datasets (Supplemental Figure S3), the negative examples were sampled uniformly for the HUANG and PAN datasets. Methods can thus primarily distinguish between positive and negative examples by node degree alone.

We further secure this finding by closer inspection of the degree ratios. Supplemental Figure S6a shows that most proteins in HUANG and PAN have either exclusively positive or negative interactions annotated (degree ratios are mainly 1 or 0). Because of the substantial data leakage (Figure 2c, Supplemental Table S2), the proportion of training proteins with a high or low degree ratio in the positive or negative parts of the test sets is very high for HUANG (87% and 86%) and PAN (92% and 90%, Supplemental Figure S7a). Suppose an algorithm correctly predicts all of these interactions because of the degree information shortcut and assigns a random label for the remaining test interactions. In that case, we expect an accuracy of 93.25% and 95.5%, respectively. This estimate is close to the prediction of most methods, including but not limited to the topology methods.

D-SCRIPT and Topsy-Turvy perform poorly due to overfitting (Supplemental Figures S8, S9). While training loss decreases and training accuracy increases, validation loss stays constant or increases, and validation accuracy decreases for all datasets. The only datasets where some learning is visible are HUANG and GUO, which is also reflected by the final reported balanced accuracy. While D-SCRIPT’s final prediction performance on the PAN dataset is far above random, inspecting the loss and accuracy patterns over the ten epochs reveals an overfitting pattern. Both models mostly predict test candidates to be non-interacting (specificity ≈1.0, see Supplemental Figure S20).

Early stopping leads to a strong improvement of D-SCRIPT and Topsy-Turvy on almost all datasets (Supplemental Figure S15). The other methods, however, mostly lose performance. Many models already reach their best performances on the validation dataset in early epochs (Supplemental Figures S8, S9), indicating that the respective datasets might not be suitable for learning as they lead to immediate overfitting.

Except for SVM-based methods, the performance of the baseline ML methods is virtually interchangeable and roughly equal to the DL methods on the larger datasets, excluding D-SCRIPT and Topsy-Turvy. The random forest-based models seem to be a powerful alternative to the deep learning models.

As expected, the performance of all methods drops significantly on RICHOUX-STRICT. As shown in Figure 2c, all datasets except for RICHOUX-STRICT include the vast majority of the proteins in both the train and test set. Consequently, RICHOUX-STRICT is less prone to training data leakage, explaining the observed results. In the presence of data leakage, robust predictions can be made based on node degree and sequence similarity, even with basic ML models. RICHOUX-STRICT’s overlap is still almost 50%, but it is free from hub biases. However, 40% of the positive and 51% of the negative test interactions still involve a protein with mainly positive or mainly negative annotations in the training set, respectively (Supplemental Figure S7). Applying the same logic as above, we expect an accuracy of 72.75%, which is the performance of most methods. SPRINT does not use node degrees for its predictions, so the number of protein pairs seen in training seems to be large enough for SPRINT to find similar subsequences for the RICHOUX-STRICT test set.

Due to memory restrictions, the harmonic function algorithm could not be run on the D-SCRIPT UNBALANCED dataset. SPRINT, both Richoux models, DeepFE, PIPR, and the random forest-based methods could handle the one to ten imbalance while the other models performed close to random (balanced accuracy around 0.5). DeepFE and PIPR held up the good performance under early stopping and D-SCRIPT and Topsy-Turvy profited strongly, showing that when overfitting is prevented and the training set is large enough, models can generalize to the test set.

For all benchmark datasets, however, the overall number of proteins is far from the real number of proteins (Supplemental Table S2). It can hence be expected that the models overfit extremely on a specific subset and will not generalize when presented with unknown proteins.

### Rewiring tests

We investigated how node degree-preserving rewiring of the positive training set affects the methods in general (Explanation 2). The differences to the results on the original datasets are shown in Figure 3a. As edges are no longer biologically meaningful, the methods can only utilize degree information shortcuts to make correct predictions in the unperturbed test sets. Suppose, in the training split, protein *p* is only involved in positive (or negative) interactions. Any PPI involving *p* in the test split will likely be predicted as positive (or negative). Sequence similarities can only help when *p*^′^ is, e.g., similar to a hub protein. Then, *p*^′^ is more likely to be a hub protein (e.g., because of a shared promiscuous domain). However, sequence similarities hinting at binding sites or interacting domains cannot be used for prediction because of the rewired training data. While a slight drop in accuracy compared to the original performance would support the validity of Explanation 2, a significant drop would indicate that the models do not learn from node degrees alone. The extent of data leakage and the distribution of node degree ratios remain comparable to the original datasets (Figures 3b, S6b, and S7b).

**Figure 3:**
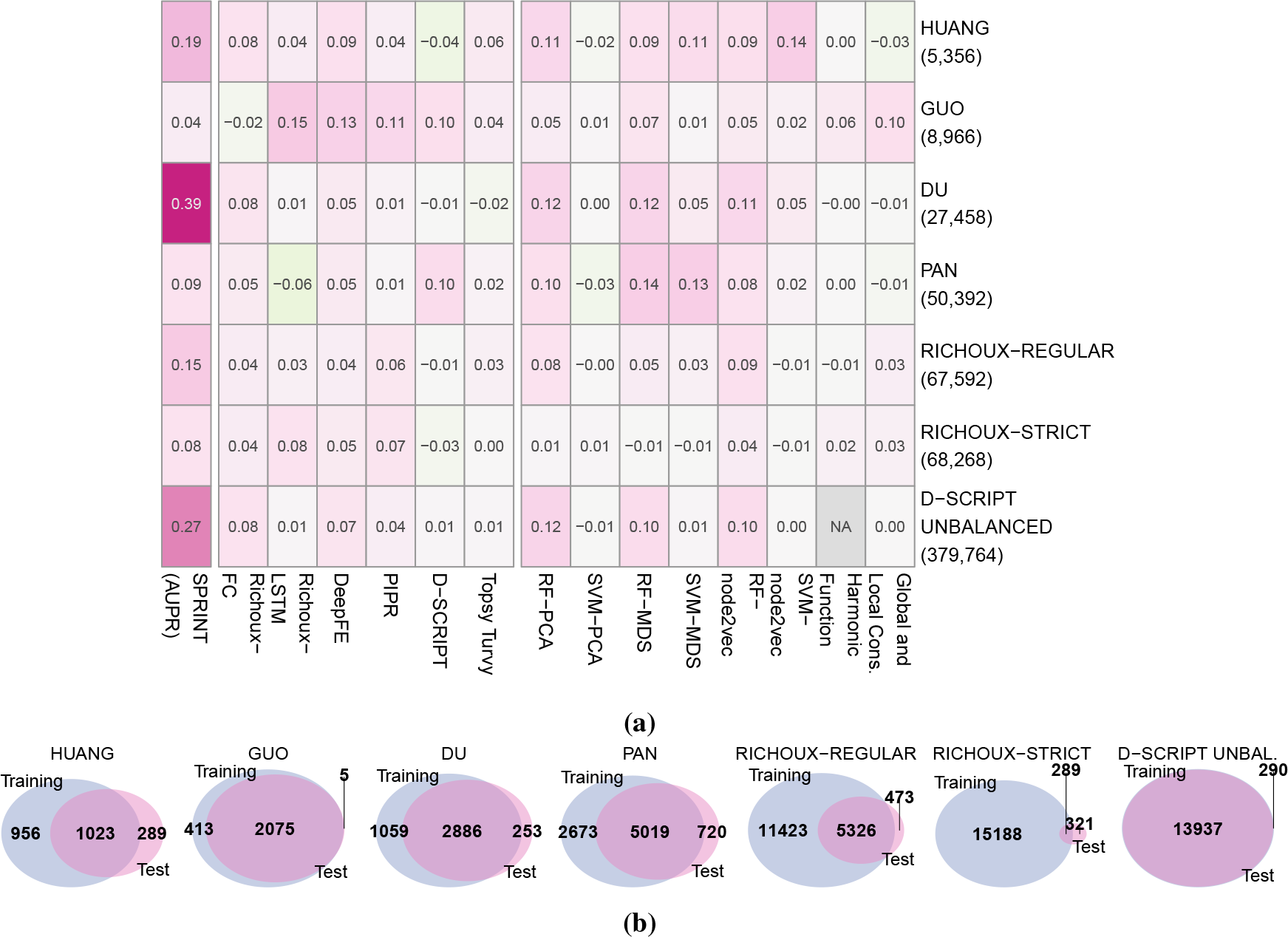
Differences between the results obtained on the original datasets and results obtained on datasets with randomly rewired positive PPIs in the train sets. **(a)** AUPR values on unmodified test sets for SPRINT, balanced accuracies for all other methods. Y-axis labels correspond to the dataset, the numbers of samples in the training data (all/restricted length) are shown in parantheses. Performances fell only slightly compared to the results on the original datasets. Very high accuracies can still be reached for HUANG and PAN. For larger datasets, basic ML models perform approximately as well as the DL models. SPRINT performs almost randomly on small datasets but still reaches accuracies around 80% on the PAN and RICHOUX datasets. **(b)** Overlap of proteins occurring in the rewired training and test sets.

Indeed, the performances fell slightly compared to the results on the original datasets for all methods. Very high accuracies can still be reached on the datasets HUANG and PAN (Supplemental Figure S14). This is in accordance with our observations from Figure 2b and our findings on the original datasets. For these two datasets, the node degree distribution of the positive PPIs (power law) is not equal to that of the negative PPIs (uniform). Additionally, the proportion of training proteins with a high or low degree ratio in the positive and negative part of the test fold is again very high (81% and 91% for HUANG and 90% and 88% for PAN, Supplemental Figure S7b). Therefore, the models can fully leverage the degree information shortcut to predict the unrewired test sets. This explanation also concurs with the observation that the sequence similarity-based baseline ML models lose more performance than the topology-based baseline methods on these datasets. In contrast to the topology-based baseline methods, sequence similarity-based baseline methods must implicitly infer the degree information shortcut.

Remarkably, D-SCRIPT gains some performance on the HUANG dataset and the loss and accuracy curves indicate some learning. Richoux-LSTM has the largest gain in performance on the PAN dataset but here, no learning can be seen and in the early stopping setting, the model from epoch 1 was taken. Generally, the random forest-based baseline models lose more accuracy points than the DL models, except on RICHOUX-REGULAR and RICHOUX-STRICT, where the performance is similar. It is possible that the DL methods are better at recognizing node degree biases compared to basic models, which need larger datasets to achieve this. This trend is best visible in the unbalanced D-SCRIPT dataset (Supplemental Figure S14).

SPRINT shows comparably poor performance on the smaller datasets but achieves an AUPR of up to 84% on large datasets. We cannot fully explain these high AUPR values. SPRINT searches for sequence similarities in potentially interacting protein pairs and does thus not benefit from node degree information. While we see a significant drop in the magnitude of the scores compared to the original datasets (Supplemental Figure S5), scores of interacting proteins are still higher than those of non-interacting proteins.

Overall, we can confirm that the methods are biased by node degree information: Balanced accuracies up to 97% can still be reached despite the rewiring of the training data.

### Partitioning tests

Running the baseline methods on the original datasets was a positive test for Explanations 2 and 3, as we expected similar performance for DL and basic ML models. Indeed, our results confirm these expectations (see Figure 2a). The partitioning tests served as a negative test for Explanations 2 and 3. If the explanations were valid, we would expect the performances to drop significantly when the models are trained on *INTRA*_0_ and tested on *INTRA*_1_. We expected this for both the DL and baseline ML models.

The results of the partitioning tests are shown in Figure 4. Notably, all training dataset sizes were approximately halved because of the partitioning strategy. While *INTER* and *INTRA*_0_ are approximately equal in size, *INTRA*_1_ is considerably smaller (see Methods for details).

**Figure 4:**
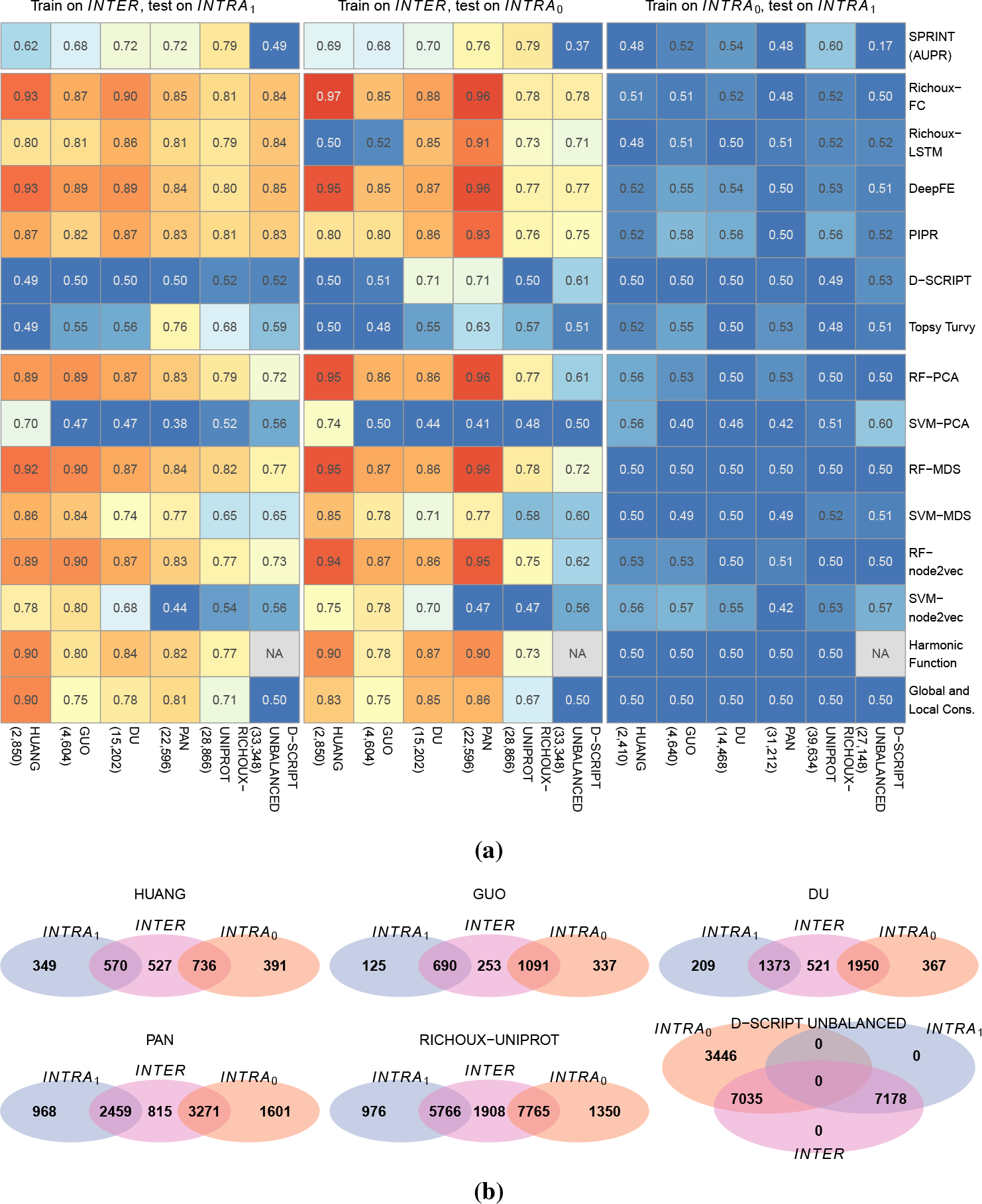
Results of partitioning tests. **(a)** AUPR values for SPRINT, accuracies for all other methods. X-axis labels correspond to the dataset, the numbers of samples in the training data are shown in parentheses. Performances drop to random for all methods when trained on *INTRA*_0_ and tested on *INTRA*_1_. Performances obtained from training on *INTER* are still excellent, especially when tested on the *INTRA*_0_ blocks of HUANG and PAN. **(b)** Overlap of proteins occurring in the different blocks of the partitions of the benchmark datasets.

Indeed, we observed random or near-to-random performances for all methods trained on *INTRA*_0_ and tested on *INTRA*_1_. The results show that when the test sets do not suffer from data leakage, the methods do not learn any higher-level features during training that they can apply to unseen data. Instead, models overfit on the interaction patterns of the training proteins. Predictions become random when the test set does not contain these proteins (or highly similar proteins). The topology-based baseline methods predict all candidates to interact, except for the unbalanced dataset, where all are predicted not to interact (see recall and specificity, Supplemental Figures S18, S20). D-SCRIPT and Topsy-Turvy profit from early stopping on the large datasets (Supplementary Figure S15). However, for both PAN and D-SCRIPT UNBALANCED, the model from the first epoch had the best performance on the validation set (Supplementary Figures S12, S13).

Overall, the performances obtained after training on *INTER* were excellent, especially when the methods were tested on the *INTRA*_0_ blocks of HUANG and PAN. Looking at the degree ratio proportions (Supplementary Figure S7), the proportion of *INTER* proteins with a low degree ratio in the negative part of the *INTRA*_0_ block is remarkably high compared to the *INTRA*_1_ block. Consequently, the models can leverage node degree information exceptionally well for the non-interactions, reflected in the high specificity achieved in this setting (Supplementary Figure S12.) Richoux-LSTM yielded random predictions for the small datasets due to overfitting, which was foreseeable when running 100 epochs on less than 4000 data points. D-SCRIPT and Topsy-Turvy also show overfitting patterns despite their objectively good performance on PAN and RICHOUX-UNIPROT (Topsy-Turvy, training on *INTER*, testing on *INTRA*_*1*_), and DU AND PAN (D-SCRIPT, training on *INTER*, testing on *INTRA*_*0*_, Supplementary Figures S12, S13).

Data leakage is highest for the D-SCRIPT UNBALANCED dataset, where all proteins of *INTRA*_1_ are contained in *INTER*_1_. This explains why all models perform better on the *INTRA*_1_ block than on the *INTRA*_0_ block. Looking at the performance developments in the early stopping setting, we can hypothesize that the models find the degree shortcut in the process of overfitting to the training set. Overall, the results obtained show that the effect of the resulting data leakage is considerable even when only one of the proteins of each training PPI occurs in the test set.

### Runtime

Runtime typically grows linearly with the size of the training dataset (Figure 5). An exception is SPRINT which always loads the preprocessed proteome before reading in the positive training PPIs. As this data contains the computed similar sub-sequences, the file is much more extensive for the human proteome compared to yeast; therefore, SPRINT takes longer on human compared to yeast datasets. Also, the topology-based baseline methods have almost constant runtime. However, the harmonic function required more than 350 Gb of memory for the D-SCRIPT UNBALANCED dataset. D-SCRIPT and Topsy-Turvy have by far the highest runtime. Our baseline ML models are the fastest methods. The fastest DL method is Richoux-FC, which only runs for 25 epochs instead of 100, like Richoux-LSTM.

**Figure 5:**
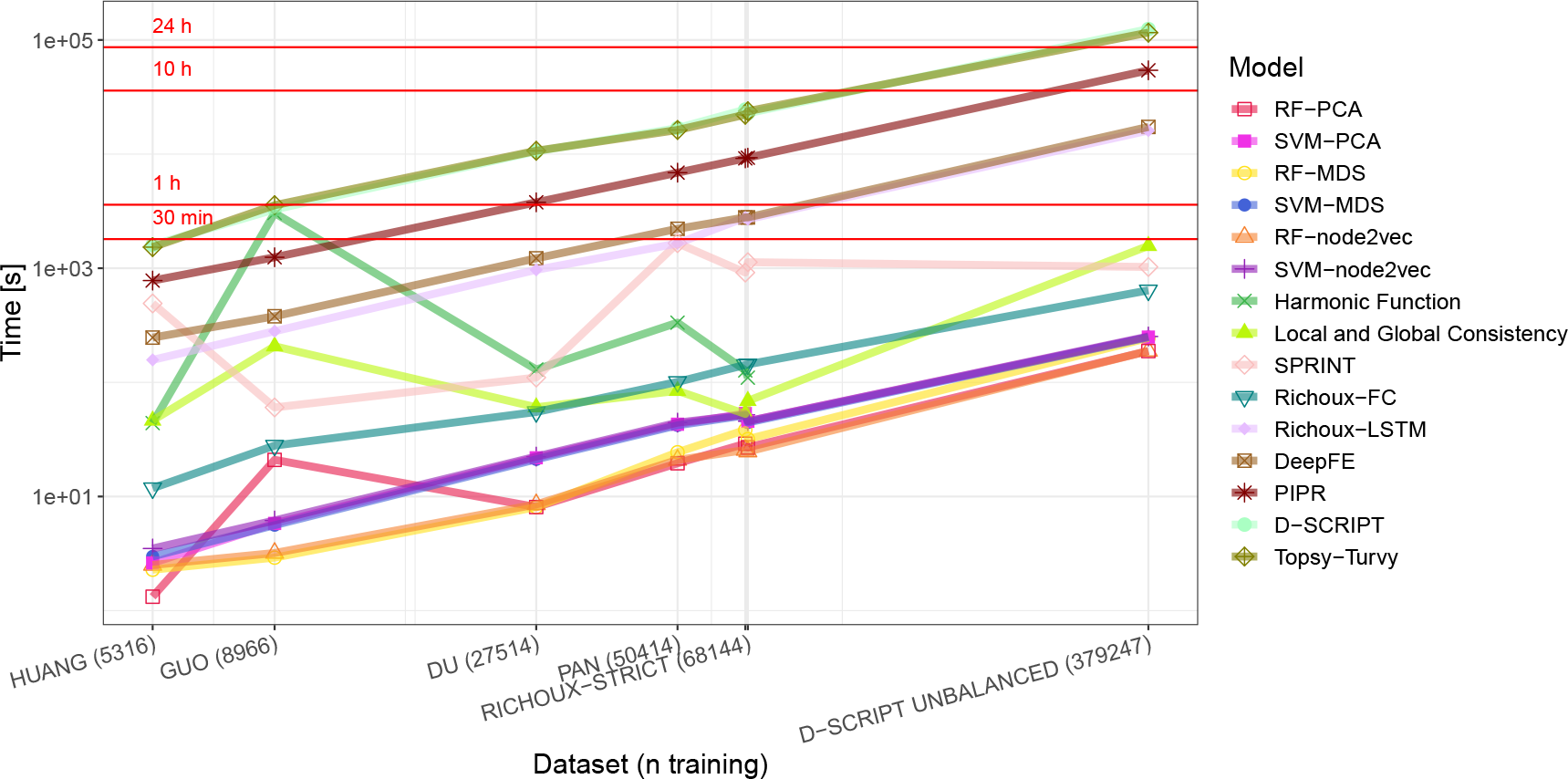
Overall runtime for training and testing on the original datasets. The label for RICHOUX-REGULAR is missing from the x-axis because it would overlap with the RICHOUX-STRICT label. Most runtimes increase linearly with size of the training dataset. D-SCRIPT and Topsy-Torvy have by far the highest runtime while the Random Forest Methods have the lowest.

**Figure 6:**
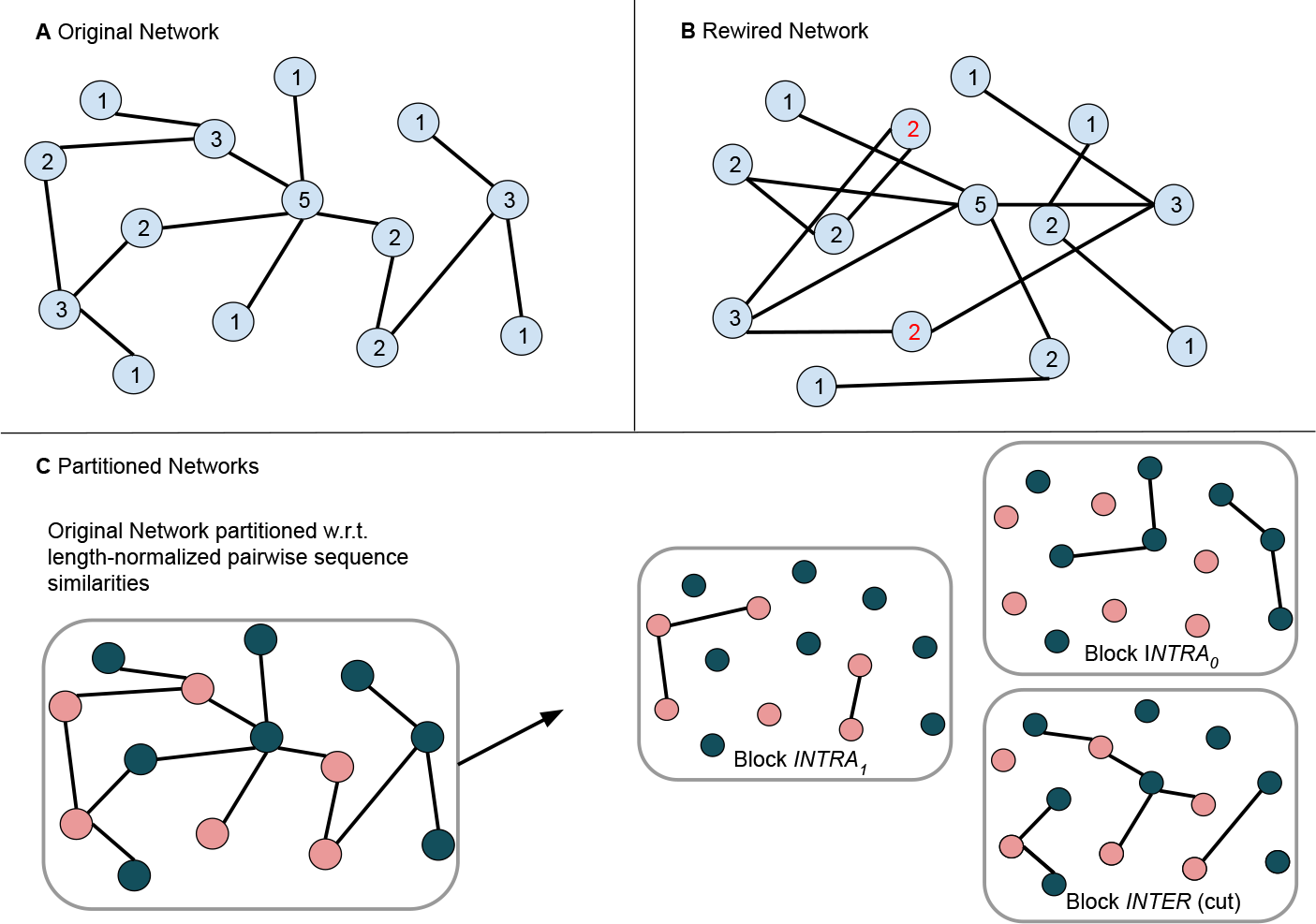
Concept of the rewiring and partitioning strategies. When the original network (A) is rewired, all node degrees stay the same in expectation but the edges are no longer meaningful (B). For the partitioning tests (C), the proteome is partitioned into two blocks (pink and green nodes) such that pairwise inter-block sequence similarity is minimized. Then, the PPIs from the original network (A) are partitioned based on which block the involved proteins are contained in (*INTRA*_0_: both proteins contained in green block, *INTRA*_1_: both proteins in contained pink block, *INTER*: proteins contained in different blocks).

Combined with the results from the previous sections, it is evident that an enormous amount of resources can be saved by using simple methods to predict PPIs like scoring algorithms, Random Forests, or multilayer perceptrons with few layers. These methods achieve similar results as the costly deep neural networks and can confidently predict PPIs using sequence similarities and learned interaction patterns for known proteins.

### Gold standard dataset

We showed that existing DL models fail to extract more complex sequence features for predicting PPIs. For the design of more advanced machine learning strategies, we provide a leakage-free human gold standard data set for training, validation, and testing (available at https://doi.org/10.6084/m9.figshare. 21591618.v3)^48^. The positive dataset was created using data from HIPPIEv2.3^49^. Negative PPIs were sampled randomly, but such that individual node degrees are preserved in expectation (Supplemental Figure S3). This dataset was split using our partitioning strategy with KaHIP, i.e., there are no overlaps between the three sets and sequence similarity is minimal. Both training and validation datasets are large enough to allow DL methods to avoid overfitting. Additionally, the sets are redundancy-reduced w.r.t. pairwise sequence similarity using CD-HIT at a 40% threshold^50^. As a result, proteins are also pairwise dissimilar within their set, such that models have to extract features beyond sequence similarity to achieve good performance.

To confirm that our gold standard data set shows the expected behavior, we evaluated all methods on it. Because SPRINT and our baseline models do not have any tunable parameters, we collapsed the training and validation set for their training. The same was done for D-SCRIPT and Topsy-Turvy because they only update their weights using the training dataset. All methods were evaluated on the test set. Indeed, performances were random for all methods (Supplemental Table S5). None of the methods could extract any higher-level features during training that could be applied to predict the test set. Topsy-Turvy was the best-performing method at an accuracy of 56%.

In the early stopping setting, we did not collapse the training and validation set but used the validation dataset to determine the best model. However, no significant changes in performance could be seen. Here, the best-performing method was D-SCRIPT with an accuracy of 55%.

## Discussion

We have conclusively shown that the problem of binary PPI prediction is not solved but wide open. Numerous publications report accuracy values between 90% and 100% and fuel a feedback loop of over-optimism. We have shown that their prediction estimates can be solely attributed to data leakage caused by random splitting into train and test sets. The datasets used in the literature cause the models to overfit based on protein homology and node degree information. More complex sequence features representing binding pockets, protein domains, or similar motifs are not extracted. Instead, methods depend on global sequence similarity and node degree.

We have reached our conclusion using three experimental settings: Firstly, we have shown that after random 80/20 splitting of the datasets, DL and baseline ML methods yield interchangeably high results on all datasets. SPRINT, the most straightforward method, performs excellent on large datasets, which shows that finding similar subsequences to predict PPIs is already sufficient to reach exceptional performance measures. When the methods cannot use information about hub proteins (RICHOUX-STRICT test set), performance drops for all methods.

Secondly, we have demonstrated that biologically meaningless edges which preserve the expected node degree do not lead to random predictions. While, for all methods, performance measures fall compared to the original datasets, accuracies up to 97% can still be reached for the DL methods and 89% for the baseline methods. Hence, the models can confidently predict PPIs from node degree information shortcuts only.

Finally, we proved that excluding training proteins from the test set and minimizing pairwise sequence similarities between training and test sets strips all methods of their predictive power. Taken together, our results show that DL methods do not learn any higher-level structural features. Conversely, we observe strongly elevated performance scores after training the methods on a data set with shared proteins.

In this study, we focused exclusively on sequence-based methods. In future work, it would be interesting to see if our findings translate to methods predicting PPIs from 3D structures. Since proteins interact in a folded state, methods using this information might extract the actual underlying patterns and find matching sites and domains. There are also algorithmic methods^51,52^ and machine learning models^5,18,41,53^ explicitly relying on phylogeny and co-evolution, whose additional value could be interesting to explore.

We would also like to point out that PPI networks neglect differences in the interactions of protein isoforms^54^, where, for instance, the absence of a binding domain will limit a protein’s set of interaction partners. Moreover, most proteins work in larger complexes, i.e., they form long-lasting interactions between two or more proteins. These complexes might then perform their functions by interacting transiently with other proteins or protein complexes. If we want to predict and understand the underlying mechanisms of PPIs, we have to consider non-binary interactions as well as the difference between transient interactions and protein complex formation.

Binary PPI predictions are already very accurate for proteins seen during training or for proteins that share similar (sub-)sequences. We have seen that high accuracies can be obtained with simple scoring methods or fast-converging ML models. We appeal to the community to first try simple methods that cost fewer resources before moving on to complex and deep model architectures. Not only are these models prone to overfitting, but they also waste an unnecessary amount of time, memory, energy, and CO2.

Notably, the sequence- and topology-based methods we have tested here are not equipped for predicting interactions in the “dark protein-protein interactome”^55^, i.e., currently understudied proteins for which no similar sequences are found in existing PPI networks. Existing methods can not adequately address this challenging scenario, and we expect that methods leveraging structural information will close this gap in the future. As impressively shown by AlphaFold2, DL models have tremendous potential. Similarly to AlphaFold1, D-SCRIPT predicts a contact-map as an intermediate step to predicting interactions. Nevertheless, we observe poor performance for D-SCRIPT and the related Topsy-Turvy, which is only partially improved by early stopping.

We speculate that extensive training data leakage has concealed the full scope of the binary PPI prediction challenge. For future ML efforts, we thus provide a large gold standard training, validation, and test set that is free from data leakage and has minimized pairwise sequence similarities. With this, we hope to kindle renewed interest in this ML challenge and motivate further progress in the refinement of existing PPI prediction networks.

## Methods

### Datasets

We tested on the seven datasets summarized in Table 1. The yeast dataset GUO^37^ contains 5594 positive PPIs from DIP with less than 40% pairwise sequence identity and 5594 negative PPIs generated from pairs of proteins appearing in the positive set, which, according to Swiss-Prot annotations, are expressed in different subcellular locations. The yeast dataset DU^12^ was generated similarly and contains 17257 positive and 48594 negative PPIs. The human dataset HUANG^40^ contains 3899 positive experimentally verified PPIs from HPRD with less than 25% pairwise sequence identity and 4262 negative PPIs, which were generated like the ones of the datasets GUO and DU. The human dataset PAN^38^ contains 36630 positive PPIs from HPRD and 36480 negative PPIs generated by combining protein pairs obtained via the approach described above with non-interacting pairs contained in the Negatome^56^. The human dataset RICHOUX-REGULAR contains positive PPIs retrieved from UniProt and negative PPIs generated by randomly pairing proteins from the positive set. Sequences were filtered to be at most 1166 amino acids long, mirror copies were added (for each PPI (*p*_1_, *p*_2_), add (*p*_2_, *p*_1_)), and the resulting dataset was split into a training (*n*_train_ = 85104), a validation (*n*_val_ = 12822), and a test fold (*n*_test_ = 12822). The human dataset RICHOUX-STRICT^2^ was constructed from the RICHOUX-UNIPROT dataset as follows: PPIs whose involved proteins appear less than 3 times were assigned to the test fold. The remainder was redistributed among the training and validation datasets. The resulting sizes of the training, validation, and test folds are, respectively, *n*_train_ = 91036, *n*_val_ = 12.506, and *n*_test_ = 720. The D-SCRIPT UNBALANCED^20^ dataset contains 43128 positive and 431379 negative PPIs, split into training (38344 positives / 383448 negatives) and test set (4794 positives / 47931 negatives). The positive PPIs are experimentally verified interactions downloaded from STRING, with lengths between 50 and 800 amino acids. Highly redundant sequences (≥40% pairwise sequence identity) were removed. Negative PPIs were generated from the positive set at a one to ten ratio to reflect that there are much more non-interacting proteins than interacting proteins.

The seven datasets were cleaned from duplicates and checked for overlaps. The training and validation folds in RICHOUX-REGULAR and RICHOUX-STRICT were joined for all analyses. All datasets except RICHOUX-REGULAR, RICHOUX-STRICT, and D-SCRIPT UNBALANCED were randomly split into a train (80%) and test (20%) set. A validation set was not needed since we omitted hyperparameter optimization. For the early stopping setting, we used 10% of the train set and a patience of 5. This set was used to determine the model with the best validation accuracy (precision for DeepFE since the method was optimized for that in the original publication), which was later used to predict the test set. After splitting, the datasets were balanced either by randomly dropping negatives or by sampling new negatives such that both proteins are already part of the dataset and that the interaction is not part of the existing positive or negative interactions (Table 2). The imbalance of the D-SCRIPT UNBALANCED dataset was maintained to test its influence on method performance.

**Table 2:** State of the benchmark datasets after cleaning and balancing: *n* denotes the overall number of samples in the datasets, i.e., the number of PPIs plus the number of randomly sampled non-edges. *n*_train_ and *n*_test_ are defined analogously for the train and test sets. The modifications were done to clean and balance the original benchmark datasets, i.e., to ensure that the number of positive PPIs (edges) equals the number of negative PPIs (non-edges) in the train and test splits of all datasets.

| Dataset | $n$ | $n_{\text{train}}$ | $n_{\text{test}}$ | Modifications |
| --- | --- | --- | --- | --- |
| GUO | 11 162 | 8966 | 2196 | 24 duplicates, sampled 35 negatives for training, dropped 37 negatives for testing |
| DU | 34 512 | 27 514 | 6998 | 2511 duplicates, dropped 23 157 negatives for training and 5680 for testing |
| HUANG | 6690 | 5316 | 1374 | 0 duplicates, dropped 619 negatives for training and 110 for testing |
| PAN | 62 962 | 50 414 | 12 548 | 55 duplicates, dropped 1678 negatives for training and 476 for testing |
| RICHOUX-REGULAR | 79 868 | 67 404 | 12 464 | 5047 duplicates, dropped 25 475 negatives for training and 342 for testing |
| RICHOUX-STRICT | 68 664 | 68 144 | 520 | 5341 duplicates, dropped 30 057 negatives for training and 200 for testing |
| D-SCRIPT UNBAL. | 426 492 | 379 247 | 47 245 | 57 duplicates, sampled 14 081 negatives for training, sampled 1660 negatives for testing |

Because of GPU restrictions, we created length-restricted versions for all datasets for D-SCRIPT and Topsy-Turvy, where each protein’s length was restricted to lie between 50 and 1000 amino acids (Table 1).

### Rewiring tests

In order to test how much the models learn from node degree only, we rewired the positive PPIs of all described training datasets such that all proteins keep their original degree in expectation. However, the edges are newly assigned, rendering them biologically meaningless. For this, we used the `expected degree graph()` function of the NetworkX 2.8 Python package. Given the list (*w*_0_, *w*_1_, …, *w*_*m*−1_) of node degrees in the original network (positive PPIs in training fold), the function constructs a graph with *m* nodes and assigns an edge between node *u* and node *v* with probability 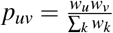. Again, all datasets were checked for duplicates and overlaps and were balanced after splitting, resulting in the counts summarized in Table 3.

**Table 3:** State of the benchmark datasets after rewiring the positive training PPIs and balancing the datasets.

| Dataset | $n$ | $n_{\text{train}}$ | $n_{\text{test}}$ | Modifications |
| --- | --- | --- | --- | --- |
| GUO | 11 256 | 8966 | 2290 | 24 duplicates, dropped 12 negatives for training, sampled 57 negatives for testing |
| DU | 34 416 | 27 458 | 6958 | 2511 duplicates, dropped 23 086 negatives for training and 5711 for testing |
| HUANG | 6722 | 5356 | 1366 | 0 duplicates, dropped 595 negatives for training and 118 for testing |
| PAN | 62 974 | 50 392 | 12 582 | 55 duplicates, dropped 1706 negatives for training and 442 for testing |
| RICHOUX-REGULAR | 80 056 | 67 592 | 12 464 | 5047 duplicates, dropped 25 382 negatives for training and 342 for testing |
| RICHOUX-STRICT | 68 788 | 68 268 | 520 | 5341 duplicates, dropped 29 996 negatives for training and 200 for testing |
| D-SCRIPT UNBAL. | 427 009 | 379 764 | 47 245 | 57 duplicates, sampled 14 832 negatives for training and 1660 for testing |

A significant drop in accuracy compared to the performance on the original dataset could indicate that the models learn from the sequence features. However, a small drop would indicate that the models mostly memorize node degrees and assign their predictions based on whether or not the protein is overall likely to interact (Explanation 2).

### Partitioning tests

To explore Explanations 2 and 3, which hypothesize that the models mostly learn from node degree information shortcuts and sequence similarities (see Introduction), we partitioned the yeast and human proteomes into two disjoint subsets *P*_0_ and *P*_1_ such that proteins from different subsets are pairwise unsimilar. For this, we first exported the yeast and human similarity networks by SIMAP2 as METIS files with length-normalized bitscore weights:

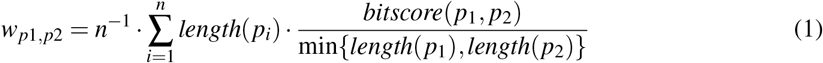

This resulted in weighted similarity networks with, respectively, 6718 nodes and 92409 edges (for the yeast proteome) and 20353 nodes and 1900490 edges (for the human proteome). In the similarity networks, bitscore edge weights increase with increasing pairwise sequence similarity.

These similarity networks were then given to the KaHIP KaFFPa algorithm (desired output partitions: 2, pre-configuration: strong), which (heuristically) solves the following problem: Given a graph *G* = (*V, E, ω*) with non-negative edge weights *ω* : *E* → ℝ_≥0_, it partitions *V* into blocks *P*_0_ and *P*_1_ such that, for all *i* ∈ {0, 1}, it holds that 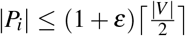(partition is almost balanced) and the total cut size 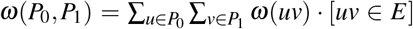 is minimized (the hyperparameter *ε* was left at the default *ε* = 0.03). For both the yeast and the human proteome, we hence obtained two disjoint subsets of proteins such that the overall pairwise sequence similarity between the subsets (sum of normalized bitscores along the cut) is minimized.

For the yeast proteome, this resulted in |*P*_0_| = 3458 and |*P*_1_|= 3260; for the human proteome, we obtained |*P*_0_|= 10481 and |*P*_1_|= 9872. Based on the partition {*P*_0_, *P*_1_ } of the human and yeast proteomes, we then partitioned each PPI dataset into three blocks *INTRA*_0_, *INTRA*_1_, and *INTER*. Each PPI (*p*_1_, *p*_2_) was assigned to one of these blocks as follows:

- We assigned (*p*_1_, *p*_2_) to the block *INTRA*_0_ if *p*_1_, *p*_2_ ∈ *P*_0_.
- We assigned (*p*_1_, *p*_2_) to the block *INTRA*_1_ if *p*_1_, *p*_2_ ∈ *P*_1_.
- We assigned (*p*_1_, *p*_2_) to the block *INTER* if *p*_1_ ∈ *P*_0_ ∧ *p*_2_ ∈ *P*_1_ or *p*_1_ ∈ *P*_1_ ∧ *p*_2_ ∈ *P*_0_.

Again, all datasets were cleaned from duplicates and balanced after partitioning. If additional negatives had to be sampled, they were sampled from the proteins of the respective block. This yielded the number of samples shown in Table 4. Methods were then either trained on block *INTRA*_0_ and tested on block *INTRA*_1_ or trained on block *INTER* and tested on the blocks *INTRA*_0_ and *INTRA*_1_. Following Explanations 2 and 3, we expected the most significant drop in accuracy compared to the original performance when training on *INTRA*_0_ and testing *INTRA*_1_. We expected a smaller drop in performance when training block *INTER* and testing on *INTRA*_0_ and *INTRA*_1_, since then, for approximately half of the test PPIs, sequence similarity information and node degrees from training are available at test time. Note that, from the three datasets published by Richoux et al.^2^, we partitioned the dataset RICHOUX-UNIPROT as it contains the largest number of unique proteins.

**Table 4:** Number of samples contained in each block after splitting the benchmark datasets according to the partitioning assignments. *n*_0_, *n*_1_, and *n*_*INTER*_ denote the numbers of positive and negative PPIs in the blocks *INTRA*_0_, *INTRA*_1_, and *INTER*. All blocks are balanced (50% interactions, 50% non-interactions).

| Dataset | $n_0$ | $n_1$ | $n_{INTER}$ | Modifications |
| --- | --- | --- | --- | --- |
| GUO | 4640 | 1722 | 4604 | $INTRA_0$: 218 additional negatives; $INTRA_1$: 86 deleted negatives; $INTER$: 307 deleted negatives |
| DU | 14468 | 4842 | 15202 | $INTRA_0$: 9686 deleted negatives; $INTRA_1$: 4707 deleted negatives; $INTER$: 14357 deleted negatives |
| HUANG | 2410 | 1426 | 2850 | $INTRA_0$: 245 additional negatives; $INTRA_1$: 66 deleted negatives; $INTER$: 544 deleted negatives |
| PAN | 31212 | 9150 | 22596 | $INTRA_0$: 180 additional negatives; $INTRA_1$: 1422 additional negatives; $INTER$: 3601 deleted negatives |
| RICHOUX-UNIPROT | 39634 | 10334 | 28866 | $INTRA_0$: 9323 additional negatives; $INTRA_1$: 4997 deleted negatives; $INTER$: 6124 deleted negatives |
| D-SCRIPT UNBAL. | 149314 | 93137 | 183414 | $INTRA_0$: 50730 additional negatives; $INTRA_1$: 15929 deleted negatives; $INTER$: 18772 deleted negatives |

### Construction of gold standard dataset

The whole human proteome was split into three parts by running KaHIP on the all-against-all sequence similarity matrix from SIMAP2 with length-normalized bitscores. When configured to output a three-way partition, KaHIP partitions the node set *V* of an edge-weighted graph *G* = (*V, E, ω*) into blocks *P*_0_, *P*_1_, and *P*_2_ such that the cut size

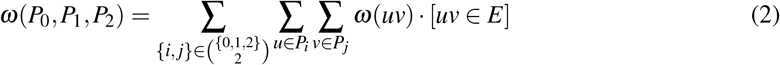

is minimized and 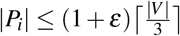 holds for all *i* 0, 1, 2. This resulted in 6987 proteins in *P*_0_, 6987 proteins in in *P*_1_, and 6379 proteins in *P*_2_. 831933 positive PPIs were downloaded from the HIPPIE database^49^ (version 2.3). Mapping all 18909 unique IDs to UniProt IDs using the UniProt mapping tool resulted in 17269 unique proteins and 689735 PPIs. The positive dataset was sorted into blocks *INTRA*_0_ (56747 PPIs), *INTRA*_1_ (164416 PPIs), and *INTRA*_2_ (52560 PPIs), where *uv*∈*INTRA*_*i*_ if and only if *u*∈*P*_*i*_ and *v*∈*P*_*i*_. Negative PPIs were sampled randomly to match the number of positives. To exclude the possibility of learning from node degrees alone, we approximately preserved the node degrees of the proteins from the positive networks *INTRA*_*i*_ in the negative networks. This was achieved by randomly sampling two distinct proteins at a time from the multiset

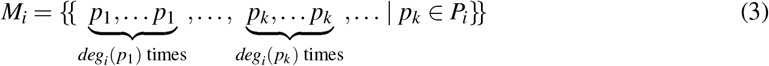

where the number of occurences of each protein *p*_*k*_ ∈ *P*_*i*_ equals its degree *deg*_*i*_(*p*_*k*_) in *INTRA*_*i*_.

Afterward, the sequences of the individual proteins in the blocks were fed to CD-HIT at a similarity threshold of 40%. Within *INTRA*_0_, CD-HIT identified 1512 redundant sequences, within *INTRA*_1_ 1680, and within *INTRA*_2_ 1465. Between *INTRA*_0_ and *INTRA*_1_, CD-HIT 2D found 3 redundant sequences, between *INTRA*_0_ and *INTRA*_2_ 20, and between *INTRA*_1_ and *INTRA*_2_ 24. These sequences were filtered out of the blocks to form redundancy-reduced datasets. The blocks were then balanced again, resulting in 59260 PPIs in *INTRA*_0_, 163192 PPIs in *INTRA*_1_, and 52048 PPIs in *INTRA*_2_. Finally, we labeled the block *INTRA*_1_ as the training dataset, the block *INTRA*_0_ as the validation dataset, and the block *INTRA*_2_ as the test dataset.

### Tested methods

The results of an extensive literature screening for high-performing PPI prediction methods can be found in Supplemental Table S1. Only twelve of 32 reviewed publications made their code available (13 methods; Richoux et al.^2^ proposed two). Eight methods were written in Python (PIPR^4^, DeepFE^13^, Richoux-FC, Richoux-LSTM^2^, D-SCRIPT^20^, Topsy-Turyv^21^, PRoBERTa^57^, and TransformerGO^46^), two in Matlab (You et al.^8^, Ding et al.^42^), one in Lua (Hashemifar et al.^18^), and two in C++ (Hamp and Rost^53^, SPRINT^36^). We excluded the methods that did not only use sequences as input (Hashemifar et al.^18^, Hamp and Rost^53^, TransformerGO^46^) and focused on DL methods with high reported accuracies, which we managed to reproduce with reasonable effort. Additionally, we included the SPRINT method as a baseline comparison since it only relies on sequence similarity for its predictions. As further similarity-based baselines, we included simple baseline ML methods (random forest, SVM), which we ran on three different encodings for the amino acid sequences (PCA, MDS, node2vec). We included two node label classification algorithms, which we ran on the line graphs of the PPI networks (harmonic function, global and local consistency) to test how much can be predicted using topology alone. Overall, we hence tested seven published methods and eight baseline ML models.

#### PIPR

PIPR^4^ encodes the two input sequences using pre-trained embeddings. These embeddings take co-occurrence similarities of amino acids and physicochemical properties into account. PIPR feeds the embeddings to a deep siamese network: the embeddings are input to two residual recurrent CNN units (convolutional layer, max pooling, bidirectional GRU with residual shortcuts, global average pooling) with shared weights. The outputs are then combined using element-wise multiplication. Finally, the network predicts binary PPIs by training a multi-layer perceptron with leaky ReLU (final activation function: softmax, loss function: binary cross-entropy, optimizer: adam with AMSgrad, learning rate= 0.001, epsilon = 1*e*^−6^). For our tests, we fixed the hyperparameters according to the results of the original publication (number of epochs: 50, batch size: 256, dimension of the convolutional and GRU layers: 25, sequence embedding cropped at position: 2000).

#### DeepFE

DeepFE^13^ pre-trained a Word2vec/Skip-gram model (size: 20, window: 4), using all Swissprot protein sequences (from 2018) as input. The model treats every amino acid as a word and, therefore, a sequence as a sentence. PPI candidate sequences are then represented using the Word2vec embeddings and are fed to a deep siamese network consisting of four units (dense layer with ReLU, batch normalization, dropout layer) with decreasing dimensionality (2048, 1024, 512, and 128). The outputs are then concatenated and put through another unit with dimensionality 8. PPI prediction is made at the final two-dimensional layer with a soft-max activation function. A stochastic gradient descent optimizer is used (learning rate: 0.01, momentum: 0.9, decay: 0.001). For our tests, we fixed the hyperparameters according to the results of the original publication (number of epochs: 45, batch size: 256, maximum protein length fed to the model: 850).

#### Richoux-FC and Richoux-LSTM

Richoux et al.^2^ represent all sequences using one-hot encoding and present two DL models: a fully connected model and a recurrent LSTM model, which we refer to as Richoux-FC and Richoux-LSTM. Richoux-FC first flattens both sequence inputs and passes them separately through two units consisting of a dense layer (dimensionality: 20, ReLU) and a batch normalization layer. The two outputs are concatenated and passed to another unit (dense layer, dimensionality: 20, ReLU). The final layer (dimensionality: 1) predicts the PPI using a sigmoid activation function. Richoux-LSTM first extracts features from the sequence via three units (convolution, pooling, batch normalization) and a final LSTM layer. The parameters are shared for these layers for the two input sequences. Then, the outputs are concatenated and passed to a dense layer (ReLU) with another batch normalization. Finally, the LSTM model predicts the PPIs using a sigmoid activation function on the final one-dimensional dense layer. Both models use the Adam optimizer with an initial learning rate of 0.001, which reduces by 10% if the loss plateaus for 5 epochs until a minimum of 0.0008. For our tests, we fixed the hyperparameters according to the results of the original publication (number of epochs for Richoux-FC: 25, number of epochs for Richoux-LSTM: 100, batch size: 256, maximum sequence length: 1166).

#### D-SCRIPT and Topsy-Turvy

Before running the D-SCRIPT model, sequences are embedded using a pre-trained language model by Bepler and Berger^58^, which transforms a sequence of length *n* to a matrix of dimension *n*×6165. In our case, we embedded the yeast and the human proteome separately. The two embeddings of a PPI between proteins of length *n* and *m* are passed separately through a projection module with shared weights, which reduces the dimensionality to *n*×100 and *m*×100, respectively. This module consists of a dense layer (dimensionality: *n*×100 and *m*×100, respectively, with ReLu activation), followed by a dropout layer. The outputs are combined by concatenating their element-wise differences and Hadamard products and used as input for the residue contact module. A dense layer (dimensionality: 50), followed by a batch normalization layer and a ReLU activation, produces a tensor of dimensionality *n*×*m*×50. D-SCRIPT passes the matrix to a convolutional layer (width: 7), followed by a batch normalization layer and a sigmoid activation function to produce a contact prediction matrix Ĉ∈ [0, 1]^*n*×*m*^. The interaction prediction module transforms this matrix into an interaction probability by subjecting it to a max-pooling operation (size: 9), a custom global pooling operation, and a custom logistic activation function. The loss function minimized during training is a weighted sum of the binary cross-entropy loss (prediction) and a contact-map magnitude loss. We fixed all hyperparameters to the ones from the publication, i.e., training for 10 epochs with a batch size of 25 and an Adam optimizer with a learning rate of 0.001.

Topsy-Turvy computes a topology score, the GLIDE score, and the D-SCRIPT prediction probability for each PPI. The GLIDE score combines a local, neighborhood-based metric (common weighted normalized score) and a global metric (diffusion state distance) to capture the network surrounding the two protein nodes, regardless of whether they are close (local metric) or not (global metric). The score is binarized at a cutoff of 92.5. A GLIDE score prediction loss extends the part of the D-SCRIPT loss responsible for interaction probability with a relative importance of 0.2. The GLIDE score prediction loss is the binary cross-entropy loss between the D-SCRIPT prediction and the binarized GLIDE score. Hyperparameters were fixed as specified in the publication, i.e., training for 10 epochs with a batch size of 25 and an Adam optimizer with a learning rate of 0.001.

#### SPRINT

Instead of finding hidden patterns via DL, SPRINT^36^ entirely relies on the hypothesis that a protein pair that is pairwise similar to an interacting protein pair is more likely to interact. Therefore, we can use it as a baseline model to see how well a dataset can be predicted if we only use sequence similarities (Explanation 3). SPRINT searches for similar subsequences between the candidate protein pair and the database of known interactions by using an approach very similar to BLAST’s hit-and-extend approach. Instead of having consecutive initial seeds of a certain length like BLAST, SPRINT also allows for gapped initial seeds to increase the number of hits. Using all hits, SPRINT calculates similarity scores and sorts the output decreasingly by the scores where a higher score represents a higher probability for interaction. Rather than choosing an arbitrary threshold to calculate accuracies, we calculated the AUC and Area Under the Precision Recall curve (AUPR) for SPRINT.

### Baseline ML models

#### Similarity-based models

We implemented additional baseline models to compare the performance of DL methods against classical ML. Note that the idea behind implementing the baseline models is not to show that classical ML models suffice for the PPI prediction task but to explain the phenomenal accuracies reported for sequence-based DL models. Because of this, we designed our baseline models such that they can only learn from sequence similarities and node degrees: the similarities are the only input features and the node degrees can be learnt implicitly during training.

As models, we trained a random forest (sklearn 1.0.2 RandomForestClassifier, 100 decision trees, six parallel jobs, a prediction greater than 0.5 is interpreted as interaction) and an SVM (sklearn SVC, RBF kernel, maximum number of iterations: 1000, a prediction *>* 0.5 is interpreted as interaction). We adopted these hyperparameters from the sklearn defaults; no tuning was done.

Proteins were represented using a similarity vector where each entry is the pairwise similarity score (bitscore) to every other protein in the human or yeast proteome (defined by Swissprot). A PPI was encoded as the concatenation of the two embeddings. An all-against-all similarity matrix for human and yeast was computed by the team of SIMAP2^39^, yielding a 6718×6718 matrix for yeast and a 20353×20353 matrix for human. Since this dimensionality is too high, we reduced it via three approaches: PCA, MDS, and node2vec, each returning 128 dimensions.

Theoretically, preprocessing the whole dataset before splitting it into train and test is considered data leakage. In this case, it is acceptable since we consider the whole universe (proteome). Introducing data leakage via joint preprocessing usually means that we have information available during testing that we would not have in a real scenario (e.g., divergence from the mean of the dataset). However, the dimensionality reductions are computed on the similarity matrix of all known human proteins; therefore, we do not introduce any dataset-specific biases. Given a new PPI, we would already possess the embeddings of its individual proteins.

PCA and MDS yielded matrices of dimensions 6718×128 and 20353×128 for the yeast and human proteomes. For the computation of the node2vec encoding, we built a weighted similarity network from the similarity matrix such that an edge between two proteins exists if and only if their bitscore is positive. Self-loops were excluded. Since SIMAP2 uses cutoffs, some of the proteins had no edges. Because the input to node2vec is just an edge list, the resulting node2vec embeddings were smaller, resulting in matrices of dimensions 6194×128 and 20210×128 for the yeast and human proteomes. Altogether, we hence tested three different embeddings as input to two ML models, yielding six baseline models.

#### Topology-based models

Both models are node label classification algorithms implemented in NetworkX 2.8. Because the PPI labels are edge labels, we first converted all PPI networks to line graphs for this task (i.e., all edges become nodes, and all nodes become edges, Supplementary Figure S1).

Zhu et al.^34^ formulate the problem of assigning labels to unlabelled nodes in a partially labeled network (our line graph) in terms of a Gaussian random field on the graph. The solution is given by the minimum harmonic energy function, which is unique and can be calculated explicitly using matrix operations. In NetworkX 2.8, the solution is calculated iteratively. On an unweighted graph, in the first iteration, node label probabilities are calculated by looking at the labels of neighboring, labeled nodes. In the next iteration, these probabilities are updated according to the label probabilities of the neighboring nodes and the node labels. This procedure is repeated *n* times, in our case, *n* = 30.

Zhou et al.^35^ slightly modified the method presented by Zhu et al., causing information to be spread symmetrically. They also introduce a parameter specifying the relative amount of information a node receives from its neighbors (here, *α* = 0.99) and the initial labels (1−*α*). A final difference is that, in contrast to the harmonic function, the label probabilities also influence the initial node labels, so outliers can potentially be corrected. Like the harmonic function, the local and global consistency algorithm is iterated *n* times, here *n* = 30.

## Code and data availability

All datasets and code are available at https://github.com/biomedbigdata/data-leakage-ppi-prediction. The gold standard dataset is available at https://doi.org/10.6084/m9.figshare.21591618.v3.

An AIMe report^59^ specifying the details of all analyses is available at https://aime-registry.org/report/VRPXym.

## Acknowledgments

We thank Prof. Dr. Thomas Rattei and his team from SIMAP2 at the University of Vienna for kindly and quickly sharing their protein similarity data. JB and ML were supported by the German Federal Ministry of Education and Research (BMBF) within the framework of the CompLS funding concept [031L0305A (DROP2AI)]. We gratefully acknowledge the support of NVIDIA Corporation with the donation of the Titan Xp GPU used for this research. ML was additionally funded by the Deutsche Forschungsgemeinschaft (DFG, German Research Foundation) [422216132]. DBB was supported by the German Federal Ministry of Education and Research (BMBF) within the framework of the CompLS funding concept [031L0309A (NetMap)].

## Author contributions

JB, DBB, and ML designed and conceived this study and drafted the manuscript. JB implemented the test protocol and carried out the analyses. ML and DBB supervised this work.

