## Supplemental Figures and Tables for "Cracking the black box of deep sequence-based protein-protein interaction prediction"

### Cracking the black box of deep sequence-based protein-protein interaction prediction - Supplemental Figures and Tables -

Judith Bernett[1,*], David B. Blumenthal[2,†], and Markus List[1,†]

[1]Chair of Experimental Bioinformatics, TUM School of Life Sciences, Technical University of Munich, Freising, Germany
[2]Biomedical Network Science Lab, Department Artificial Intelligence in Biomedical Engineering, Friedrich-Alexander-Universität Erlangen-Nürnberg, Erlangen, Germany
[†]These authors contributed equally.

**Keywords:** PPI prediction, Data Leakage, Deep Learning

Table S1: Literature overview. Datasets: ∗ GUO (yeast), ○ PAN (human), † MARTIN (H. pylori), • Cross-species DIP dataset (E. coli: E, H. sapiens: H, C. elegans: C, M. musculus: M, H. pylori: P, D. melanogaster: D), ⋄ HUANG (human), ⊗ DU (yeast). Citations: Number of Citations reported on Google Scholar on 25.05.23. **Code available.**

| Paper | Method | Input | Reported accuracies | Reported on | Citations |
|---|---|---|---|---|---|
| 1. Guo et al. (2008)[1] | AC encoding + SVM | Sequence only. | 0.87∗, other own datasets: 0.58-0.86 | 2/5 random test | 667 |
| 2. Guo et al. (2010)[2] | SVM | Sequence only. | 0.89∗, 0.91○, 0.93 E/ 0.90 D/ 0.978 C •, other: 0.94 | 2/5 random test | 69 |
| 3. Pan et al. (2010)[3] | CT encoding + hierarchical LDA-RF | Sequence only. | 0.98○ | Mean 5CV | 176 |
| 4. Sun et al. (2017)[4] | AC encoding + stacked autoencoder | Sequence only. | 0.97○, 0.5†, 0.96 E/ 0.98 D/ 0.97 C •, other: 0.9-0.99 | ca. 1/10 random test | 317 |
| 5. **Chen et al. (2019, PIPR)[5]** | Embeddings, siamese RCNN, MLP | Sequence only. | 0.97∗ | Mean 5CV | 157 |
| 6. Wang et al. (2019)[6] | PSSM, CNN with FSRF | PSSM (PSI-BLAST) | 0.98∗, 0.89† | Mean 5CV | 77 |
| 7. Xu et al. (2020)[7] | Feature extraction: physicochem., graph energy, contact graph energy, dipeptide composition. PCA, WSRC classifier | Sequence only. | 1.0∗, 0.97†, 0.99⋄ | Mean 5CV | 9 |
| 8. Wang et al. (2017)[8] | LCTD encoding, NN | Sequence only. | 0.87†, 0.95 E/ 0.93 C/ 0.94 H/ 0.93 M •, 0.93⊗ | Mean 5CV | 39 |
| 9. **You et al. (2015)[9]** | MLD feature representation + RF | Sequence only. | 0.95∗, 0.88†, 0.89 E/ 0.88 C/ 0.94 H/ 0.92 M/ 0.91 P • | Mean 5CV | 160 |
| 10. Hu et al. (2015)[10] | frequently occurring variable-length sequence segments, patterns sign. compared to background, weight, probability for interaction | Sequence only. | 0.62 ∗ (AUC), 0.68 ○ (AUC) | Mean 5CV | 23 |
| 11. Du et al. (2017)[11] | Feature extraction, Siamese DNN | Sequence only. | 0.86†, 0.92 E/ 0.95 C/ 0.94 H/ 0.91 M •, 0.98⋄, 0.93⊗, other: 0.79-0.91 | 1/4 random test | 172 |

Table S1: Literature overview. Datasets: ∗ GUO (yeast), ○ PAN (human), † MARTIN (H. pylori), • Cross-species DIP dataset (E. coli: E, H. sapiens: H, C. elegans: C, M. musculus: M, H. pylori: P, D. melanogaster: D), ⋄ HUANG (human), ⊗ DU (yeast). Citations: Number of Citations reported on Google Scholar on 25.05.23. **Code available.**

| | | | | | |
|---|---|---|---|---|---|
| 12. **Yao et al. (2019, DeepFE)**[12] | Res2Vec encoding + siamese network | Sequence only. | 0.95∗, 1 E/ 1 C/ 1 H/ 1 M •, 0.99⋄, other: 0.73-0.94 | 2/5 random test | 51 |
| 13. Jha & Saha (2020)[13] | ResNet50 on 3D data, LSTM + Encoder/Decoder on sequence AC/CT embedding | Sequence + 3D structure | 0.97○, 0.94⊗ | Mean 3CV | 15 |
| 14. Saha et al. (2014)[14] | Ensemble Learning: SVM, RF, decision tree, naive Bayes | Overlap of GO graphs, interacting domains, paralogous verification method. | own datasets 0.68-0.91 | Mean 10CV | 54 |
| 15. Chen et al. (2019)[15] | AC encoding, GO LCA clustering, functional similarity graph, Resniks measure on GO + Ensemble learner: RF, naive Bayes, NN, KNN. SVM for classification | Sequence, GO terms, topological features | 0.84∗, 0.98 H/ 0.95 E/ 0.98 D/ 0.99 C •, 0.92⊗, other: 0.78-0.94 | Mean 10CV | 52 |
| 16. Zhao et al. (2020)[16] | sequence, GO: Bert + BIGRU, Inception CNN, Attention, GAP, GMP | Sequence + GO terms | 0.97∗, 0.90⊗, other: 0.88-0.96 | Mean 10CV | 11 |
| 17. **Hashemifar et al. (2018)**[17] | PSSM, siamese-like CNN, random projection module | PSSM (PSI-BLAST) | 0.95∗, 0.96 H/ 0.97 E/ 0.96 C/ 0.96 M • | Mean 10CV and 5CV | 248 |
| 18. Maetschke et al. (2021)[18] | Different encodings of GO terms + RF | GO terms. | 0.9∗, other: 0.73-0.93 | Mean 10CV | 85 |
| 19. Shen et al. (2007)[19] | CT encoding, SVM | Sequence only. | own dataset 0.84 | ca. 1/100, repeated 5 times | 1028 |
| 20. Mahapatra et al. (2020)[20] | AAC encoding + PCA/PSO, SVM | Sequence only. | own dataset: 0.99 | 2/5 random test | 3 |
| 21. Wang et al. (2018)[21] | PSSM, PCA, rotation forest | PSSM (PSI-BLAST) | 0.97∗, 0.88†, 0.92 H/ 1.0 E/ 0.91 C/ 0.91 M • | Mean 5CV | 35 |

Table S1: Literature overview. Datasets: ∗ GUO (yeast), ○ PAN (human), † MARTIN (H. pylori), • Cross-species DIP dataset (E. coli: E, H. sapiens: H, C. elegans: C, M. musculus: M, H. pylori: P, D. melanogaster: D), ◇ HUANG (human), ⊗ DU (yeast). Citations: Number of Citations reported on Google Scholar on 25.05.23. **Code available.**

| | | | | | |
|---|---|---|---|---|---|
| 22. **Hamp & Rost (2015)**[22] | Evolutionary profile kernel, k-mers + SVM | Sequence + PredictProtein. | own datasets: 0.67-0.87 | Mean 10CV | 107 |
| 23. **Li & Ilie (SPRINT, 2017)**[23] | spaced seeds, hit-and-extend, scores for similar subsequences | Sequence only. | own datasets: 0.74-0.93, other: 0.61-0.82 | Mean 10 CV | 65 |
| 24. **Ieremie et al. (TransformerGO, 2022)**[24] | GO-sets, graph embeddings (node2vec), Transformer | GO terms. | other: 0.91-0.97 | 2/5 random test | 11 |
| 25. **Ding et al. (2016)**[25] | Multivariate MI feature representation, Moreau-Broto Autocorrelation + RF | Sequence only. | 0.95∗, 0.88 †, 0.94 H/ 0.93 E/ 0.92 C/ 0.96 M •, 0.98 ◇ | Mean 10CV | 139 |
| 26. **Richoux et al. (FC, LSTM, 2019)**[26] | Fully connected and recurrent deep models | Sequence only. | own datasets: 0.76-0.9 | ca. 12% random and ca. 0.6% strict test set | 24 |
| 27. **Sledzieski et al. (D-SCRIPT, 2021)**[27] | Pre-trained protein-wise embedding, fully connected projection module, convolutional contact module | Sequence only. | own datasets: 0.405-0.580 auPR | Mean 5CV, cross-species test | 39 |
| 28.**Singh et al. (Topsy-Turvy, 2022)**[28] | D-SCRIPT + GLIDE score for network topology | Sequence only. | own datasets: 0.533-0.824 auPR | Mean CV, cross-species test | 6 |
| 29. **Nambiar et al. (PRoBERTa, 2020)**[29] | Task-agnostic transformer for protein sequence encoding, fine-tuning for PPI prediction (concatenated sequence) and family classification | Sequence only. | own datasets: 0.79-0.89 | 10% random test | 77 |
| 30. Pazos et al. (Mirrortree, 2001)[30] | MSA of protein orthologs for both proteins, correlate evolutionary distance matrices | Sequence, MSAs. | None, just correlations | - | 648 |

Table S1: Literature overview. Datasets: ∗ GUO (yeast), ○ PAN (human), † MARTIN (H. pylori), ● Cross-species DIP dataset (E. coli: E, H. sapiens: H, C. elegans: C, M. musculus: M, H. pylori: P, D. melanogaster: D), ◇ HUANG (human), ⊗ DU (yeast). Citations: Number of Citations reported on Google Scholar on 25.05.23. **Code available.**

| | | | | | |
|---|---|---|---|---|---|
| 31. Ochoa et al. (p-Mirrortree, 2015)[31] | Computes p-values for mirrortree scores from a background distribution | Sequence, MSAs. | own datasets: ca. 0.15 PPV, ca. 0.14 F1 | whole dataset | 27 |
| 32. Humphreys et al. (2021)[32] | Pipeline: MSAs of yeast orthologs, contact probabilities with RoseTTAFold, refinement with AlphaFold | Sequence, MSAs. | Comparison to PDB, detailed analysis of single structures | - | 231 |

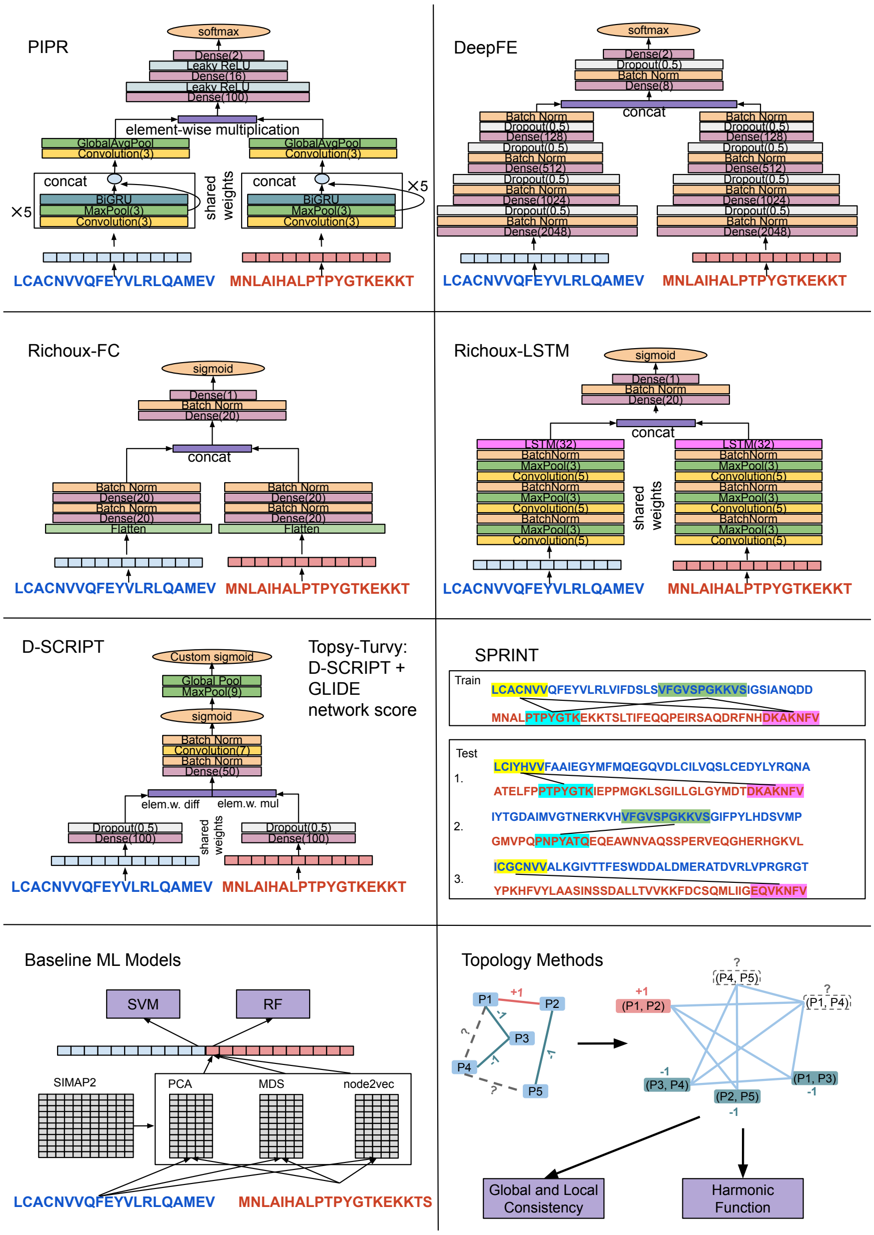


Figure S1: Overview of all tested methods. The DL methods all propose siamese network architectures where the sequences are first processed individually (shared weights for PIPR, Richoux-LSTM, and D-SCRIPT). The output is then combined and processed by fully connected layers. Predictions are made by a final softmax (PIPR, DeepFE) or sigmoid (Richoux-FC, Richoux-LSTM, D-SCRIPT) activation function. SPRINT examines candidate PPIs for subsequences similar to subsequences of known interacting proteins. Scores are calculated from pairs of subsequences, indicated by black lines. Our baseline models embed proteins using a dimensionality-reduced (PCA / MDS / node2vec) sequence-similarity vector (bitscores computed by SIMAP2). The embeddings are concatenated for a PPI and fed to either a random forest or a support vector machine. The topology methods work on the line graphs of the PPI networks and infer the missing node labels of the test set.

| Dataset | Unique Whole Data | Overlap Whole Data/Train | Overlap Train/Test | % of Test Set |
|---|---|---|---|---|
| HUANG | 2,571 | 2,354 | 1,086 | 83.3% |
| HUANG$_{LR}$ | 2,053 | 1,883 | 831 | 83.0% |
| GUO | 2,497 | 2,494 | 2,089 | 99.9% |
| GUO$_{LR}$ | 2,209 | 2,207 | 1,776 | 99.9% |
| DU | 4,417 | 4,258 | 2,969 | 94.9% |
| DU$_{LR}$ | 3,977 | 3,813 | 2,566 | 94.0% |
| PAN | 9,122 | 8,631 | 5,335 | 91.6% |
| PAN$_{LR}$ | 7,587 | 7,166 | 4,293 | 91.1% |
| D-SCRIPT UNBAL. | 14,227 | 14,227 | 13,937 | 100% |
| D-SCRIPT UNBAL.$_{LR}$ | 14,214 | 14,214 | 13,924 | 100% |
| RICHOUX-REGULAR | 17,245 | 16,847 | 5,398 | 93.1% |
| RICHOUX-REGULAR$_{LR}$ | 16,640 | 16,261 | 5,190 | 93.2% |
| RICHOUX-STRICT | 15,831 | 15,512 | 291 | 47.7% |
| RICHOUX-STRICT$_{LR}$ | 15,318 | 15,042 | 256 | 48.1% |

Table S2: Number of unique proteins occurring in the original datasets, number of unique proteins occurring in the training set, overlap of unique proteins occurring in both the train and the test set, and proportion of the overlap w.r.t the test set. Numbers are displayed for the normal and length-restricted (LR) datasets.

| Dataset | Unique Whole Data | Overlap Whole Data/Train | Overlap Train/Test | % of Test set |
|---|---|---|---|---|
| HUANG | 2,268 | 1,979 | 1,023 | 78.0% |
| HUANG$_{LR}$ | 1,835 | 1,616 | 790 | 78.3% |
| GUO | 2,493 | 2,488 | 2,075 | 99.8% |
| GUO$_{LR}$ | 2,206 | 2,197 | 1,746 | 99.5% |
| DU | 4,198 | 3,945 | 2,886 | 91.9% |
| DU$_{LR}$ | 3,783 | 3,540 | 2,476 | 91.1% |
| PAN | 8,412 | 7,692 | 5,019 | 87.5% |
| PAN$_{LR}$ | 7,030 | 6,424 | 4,036 | 86.9% |
| D-SCRIPT UNBAL. | 14,227 | 14,227 | 13,937 | 100% |
| D-SCRIPT UNBAL.$_{LR}$ | 14,214 | 14,214 | 13,923 | 100% |
| RICHOUX-REGULAR | 17,222 | 16,749 | 5,326 | 91.8% |
| RICHOUX-REGULAR$_{LR}$ | 16,634 | 16,185 | 5,124 | 91.9% |
| RICHOUX-STRICT | 15,798 | 15,477 | 289 | 47.4% |
| RICHOUX-STRICT$_{LR}$ | 15,282 | 15,008 | 252 | 47.9% |

Table S3: Number of unique proteins occurring in the rewired datasets, number of unique proteins occurring in the training set, overlap of unique proteins occurring in both the train and the test set, and proportion of the overlap w.r.t the test set. Numbers are displayed for the normal and length-restricted (LR) datasets.

| Dataset | Unique Whole Data | Whole Data ∩ Train | Train ∩ Test | % of Test Set |
|---|---|---|---|---|
| HUANG $INTER \rightarrow INTRA_0$ | 2,224 | 1,833 | 736 | 65.3% |
| HUANG$_{LR}$ $INTER \rightarrow INTRA_0$ | 1,728 | 1,442 | 542 | 65.5% |
| HUANG $INTER \rightarrow INTRA_1$ | 2,182 | 1,833 | 570 | 62.0% |
| HUANG$_{LR}$ $INTER \rightarrow INTRA_1$ | 1,774 | 1,442 | 479 | 59.1% |
| HUANG $INTRA_0 \rightarrow INTRA_1$ | 2,046 | 1,127 | 0 | 0.0% |
| HUANG$_{LR}$ $INTRA_0 \rightarrow INTRA_1$ | 1,639 | 828 | 0 | 0.0% |
| GUO $INTER \rightarrow INTRA_0$ | 2,371 | 2,034 | 1,091 | 76.4% |
| GUO$_{LR}$ $INTER \rightarrow INTRA_0$ | 2,052 | 1,752 | 915 | 75.3% |
| GUO $INTER \rightarrow INTRA_1$ | 2,159 | 2,034 | 690 | 84.7% |
| GUO$_{LR}$ $INTER \rightarrow INTRA_1$ | 1,877 | 1,752 | 611 | 83.0% |
| GUO $INTRA_0 \rightarrow INTRA_1$ | 2,243 | 1,428 | 0 | 0.0% |
| GUO$_{LR}$ $INTRA_0 \rightarrow INTRA_1$ | 1,951 | 1,215 | 0 | 0.0% |
| DU $INTER \rightarrow INTRA_0$ | 4,211 | 3,844 | 1,950 | 84.2% |
| DU$_{LR}$ $INTER \rightarrow INTRA_0$ | 3,753 | 3,403 | 1,668 | 82.7% |
| DU $INTER \rightarrow INTRA_1$ | 4,053 | 3,844 | 1,373 | 86.8% |
| DU$_{LR}$ $INTER \rightarrow INTRA_1$ | 3,632 | 3,403 | 1,239 | 84.4% |
| DU $INTRA_0 \rightarrow INTRA_1$ | 3,899 | 2,317 | 0 | 0.0% |
| DU$_{LR}$ $INTRA_0 \rightarrow INTRA_1$ | 3,486 | 2,018 | 0 | 0.0% |
| PAN $INTER \rightarrow INTRA_0$ | 8,146 | 6,545 | 3,271 | 67.1% |
| PAN$_{LR}$ $INTER \rightarrow INTRA_0$ | 6,652 | 5,370 | 2,610 | 67.1% |
| PAN $INTER \rightarrow INTRA_1$ | 7,513 | 6,545 | 2,459 | 71.8% |
| PAN$_{LR}$ $INTER \rightarrow INTRA_1$ | 6,362 | 5,370 | 1,990 | 66.7% |
| PAN $INTRA_0 \rightarrow INTRA_1$ | 8,299 | 4,872 | 0 | 0.0% |
| PAN $INTRA_0 \rightarrow INTRA_1$ | 6,874 | 3,892 | 0 | 0.0% |
| D-SCRIPT UNBAL. $INTER \rightarrow INTRA_0$ | 17,659 | 14,213 | 7,035 | 67.1% |
| D-SCRIPT UNBAL.$_{LR}$ $INTER \rightarrow INTRA_0$ | 16,015 | 14,200 | 7,023 | 79.5% |
| D-SCRIPT UNBAL. $INTER \rightarrow INTRA_1$ | 14,213 | 14,213 | 7,178 | 100% |
| D-SCRIPT UNBAL.$_{LR}$ $INTER \rightarrow INTRA_1$ | 14,200 | 14,200 | 7,177 | 100% |
| D-SCRIPT UNBAL. $INTRA_0 \rightarrow INTRA_1$ | 17,659 | 10,481 | 0 | 0.0% |
| D-SCRIPT UNBAL.$_{LR}$ $INTRA_0 \rightarrow INTRA_1$ | 16,015 | 8,838 | 0 | 0.0% |
| RICHOUX-UNIPROT $INTER \rightarrow INTRA_0$ | 16,789 | 15,439 | 7,765 | 85.2% |
| RICHOUX-UNIPROT$_{LR}$ $INTER \rightarrow INTRA_0$ | 15,985 | 14,646 | 7,287 | 84.5% |
| RICHOUX-UNIPROT $INTER \rightarrow INTRA_1$ | 16,415 | 15,439 | 5,766 | 85.5% |
| RICHOUX-UNIPROT$_{LR}$ $INTER \rightarrow INTRA_1$ | 15,645 | 14,646 | 5,459 | 84.5% |
| RICHOUX-UNIPROT $INTRA_0 \rightarrow INTRA_1$ | 15,857 | 9,115 | 0 | 0.0% |
| RICHOUX-UNIPROT$_{LR}$ $INTRA_0 \rightarrow INTRA_1$ | 15,084 | 8,626 | 0 | 0.0% |

Table S4: Number of unique proteins occurring in the partitioned datasets, number of unique proteins occurring in the training set, overlap of unique proteins occurring in both the train and the test set, and proportion of the overlap w.r.t the test set. Numbers are displayed for the normal and length-restricted (LR) datasets.

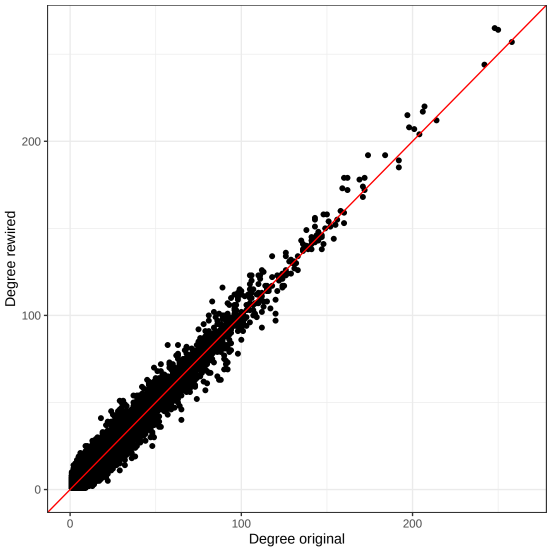


Figure S2: Node degrees of the proteins in the original vs. rewired networks. This serves merely as a sanity check to see that indeed, the rewiring preserved the node degrees for all proteins in expectation.

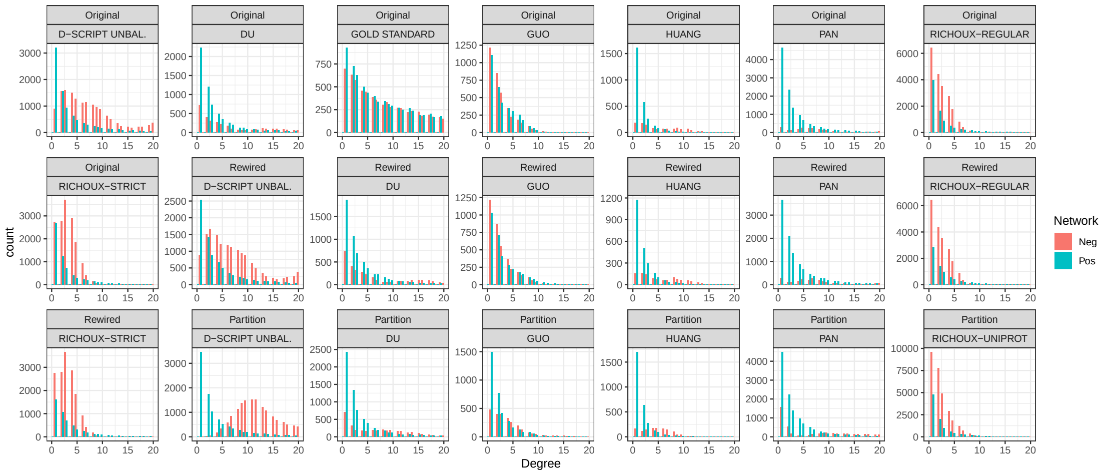


Figure S3: Comparison of the node degrees of the positive vs. negative networks for all datasets. The distributions follow the same trend for all datasets except for HUANG and PAN. For these two, negative PPIs were sampled uniformly while the positive PPI distribution follows the power law.

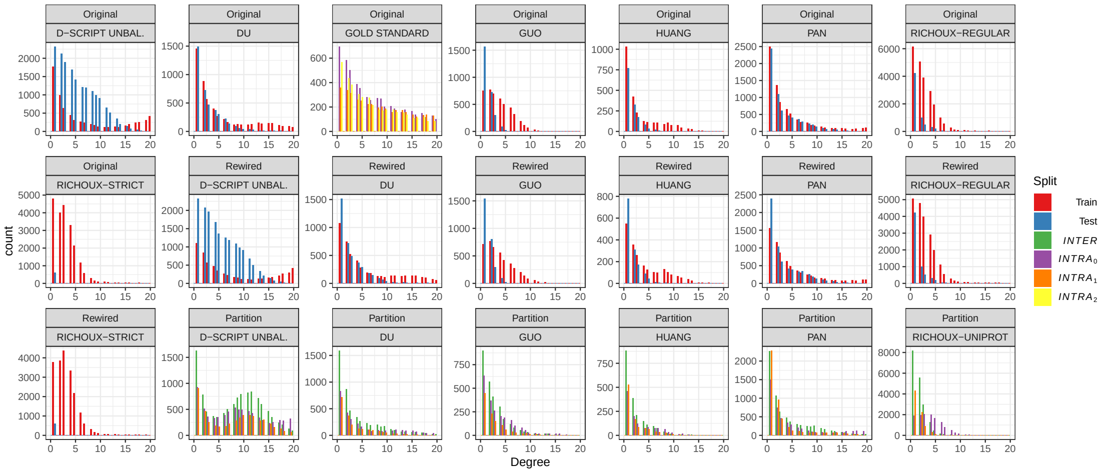


Figure S4: Comparison of node degrees between training and test sets (original/rewired) and block $INTRA_0$ vs. $INTRA_1$ vs. $INTER$. The distributions all follow approximately the same trends.

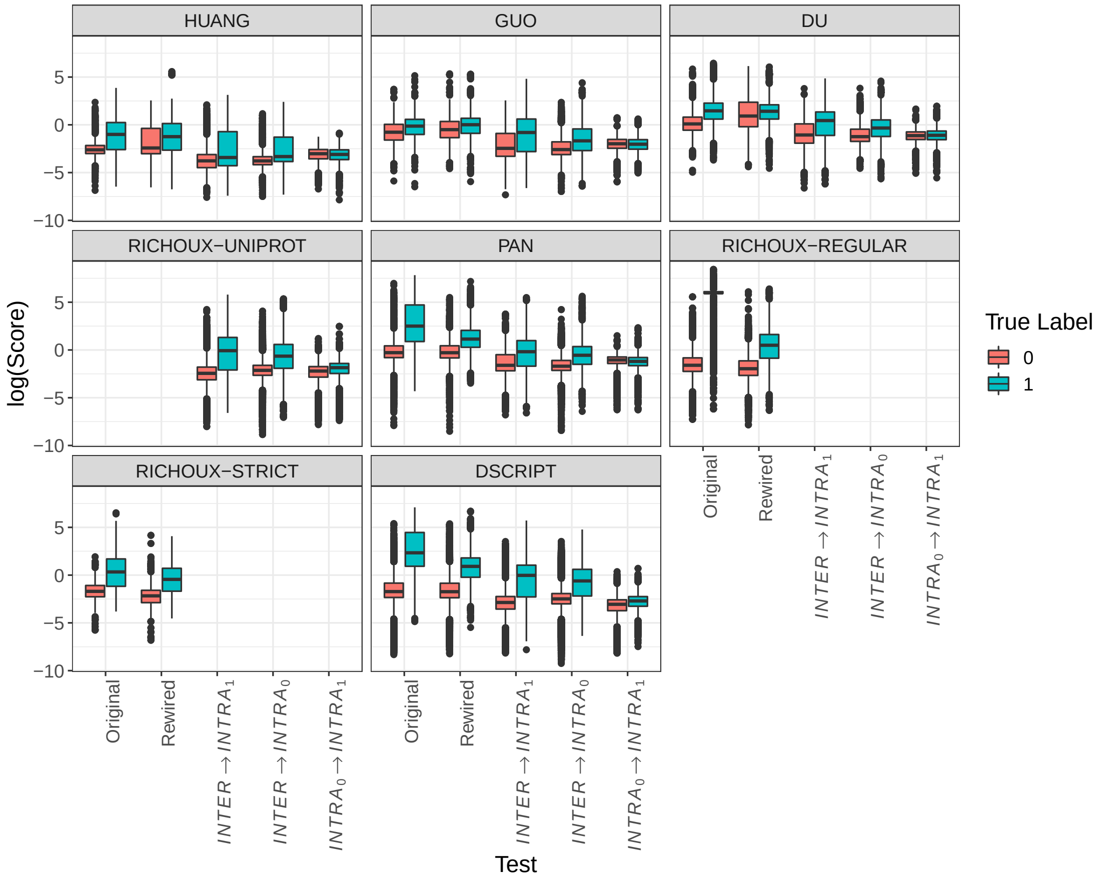


Figure S5: Distribution of SPRINT scores. For almost all original datasets, scores assigned to positive PPIs are significantly higher than scores assigned to negative examples. This difference drops strongly for the rewired test sets and the $INTER \rightarrow INTRA_0$/ $INTRA_1$ prediction (the y-axis is logged). For $INTRA_0 \rightarrow INTRA_1$, no difference between scores for positive and negative samples can be seen, resulting in random performance measures.

| Loc./ Glob. Cons. | D-SCRIPT | SVM-MDS | RF-node2vec | RF-MDS | SVM-PCA | Richoux-LSTM | SPRINT | SVM-node2vec | Deep FE | RF-PCA | PIPR | Richoux-FC | Topsy-Turvy |
|---|---|---|---|---|---|---|---|---|---|---|---|---|---|
| 0.50 | 0.50 | 0.50 | 0.51 | 0.51 | 0.51 | 0.52 | 0.52 | 0.52 | 0.52 | 0.52 | 0.52 | 0.52 | 0.56 |

Table S5: Accuracies of all methods evaluated on the Gold Standard test set. Richoux-LSTM, DeepFE, Richoux-FC, and PIPR were trained with the training set and validated on the validation set. The other methods do not have tunable hyperparameters and hence were trained on the collapsed training and validation set.

| Loc./ Glob. Cons. | Deep FE | SVM-MDS | RF-node2vec | RF-MDS | SVM-PCA | SPRINT | SVM-node2vec | RF-PCA | Richoux-LSTM | PIPR | Richoux-FC | Topsy-Turvy | D-SCRIPT |
|---|---|---|---|---|---|---|---|---|---|---|---|---|---|
| 0.50 | 0.50 | 0.50 | 0.51 | 0.51 | 0.51 | 0.52 | 0.52 | 0.52 | 0.52 | 0.52 | 0.53 | 0.53 | 0.55 |

Table S6: Accuracies of all methods evaluated on the Gold Standard test set after early stopping.

Table S7: Epochs in which early stopping was triggered.

| Model | Dataset | Original | Rewired | Partition |
|---|---|---|---|---|
| DeepFE | D-SCRIPT UNBAL. | 15 | 16 | $INTER \rightarrow INTRA_0$: 7; $INTER \rightarrow INTRA_1$: 9; $INTRA_0 \rightarrow INTRA_1$: 11 |
| DeepFE | DU | 1 | 1 | $INTER \rightarrow INTRA_0$: 1; $INTER \rightarrow INTRA_1$: 1; INTRA0->INTRA1: 2 |
| DeepFE | GUO | 2 | 1 | $INTER \rightarrow INTRA_0$: 2; $INTER \rightarrow INTRA_1$: 3; INTRA0->INTRA1: 3 |
| DeepFE | HUANG | 5 | 12 | $INTER \rightarrow INTRA_0$: 1; $INTER \rightarrow INTRA_1$: 2; INTRA0->INTRA1: 3 |
| DeepFE | PAN | 1 | 7 | $INTER \rightarrow INTRA_0$: 2; $INTER \rightarrow INTRA_1$: 2; INTRA0->INTRA1: 1 |
| DeepFE | RICHOUX | | | $INTER \rightarrow INTRA_0$: 2; $INTER \rightarrow INTRA_1$: 1; INTRA0->INTRA1: 1 |
| DeepFE | RICHOUX-REGULAR | 1 | 1 | |
| DeepFE | RICHOUX-STRICT | 1 | 1 | |
| PIPR | D-SCRIPT UNBAL. | 10 | 1 | $INTER \rightarrow INTRA_0$: 1; $INTER \rightarrow INTRA_1$: 2; INTRA0->INTRA1: 1 |
| PIPR | DU | 9 | 4 | $INTER \rightarrow INTRA_0$: 6; $INTER \rightarrow INTRA_1$: 16; INTRA0->INTRA1: 10 |
| PIPR | GUO | 1 | 1 | $INTER \rightarrow INTRA_0$: 12; $INTER \rightarrow INTRA_1$: 15; INTRA0->INTRA1: 6 |
| PIPR | HUANG | 28 | 1 | $INTER \rightarrow INTRA_0$: 6; $INTER \rightarrow INTRA_1$: 1; INTRA0->INTRA1: 14 |
| PIPR | PAN | 3 | 8 | $INTER \rightarrow INTRA_0$: 22; $INTER \rightarrow INTRA_1$: 9; INTRA0->INTRA1: 7 |
| PIPR | RICHOUX | | | $INTER \rightarrow INTRA_0$: 12; $INTER \rightarrow INTRA_1$: 8; INTRA0->INTRA1: 6 |
| PIPR | RICHOUX-REGULAR | 13 | 4 | |

Table S7: Epochs in which early stopping was triggered.

| | | | | |
|---|---|---|---|---|
| PIPR | RICHOUX-STRICT | 6 | 7 | |
| Richoux FC | D-SCRIPT UNBAL. | 1 | 1 | $INTER \rightarrow INTRA_0$: 1; $INTER \rightarrow INTRA_1$: 2; INTRA0->INTRA1: 2 |
| Richoux FC | DU | 1 | 1 | $INTER \rightarrow INTRA_0$: 1; $INTER \rightarrow INTRA_1$: 3; INTRA0->INTRA1: 1 |
| Richoux FC | GUO | 1 | 1 | $INTER \rightarrow INTRA_0$: 1; $INTER \rightarrow INTRA_1$: 25; INTRA0->INTRA1: 1 |
| Richoux FC | HUANG | 1 | 1 | $INTER \rightarrow INTRA_0$: 1; $INTER \rightarrow INTRA_1$: 1; INTRA0->INTRA1: 3 |
| Richoux FC | PAN | 16 | 1 | $INTER \rightarrow INTRA_0$: 1; $INTER \rightarrow INTRA_1$: 25; INTRA0->INTRA1: 25 |
| Richoux FC | RICHOUX | | | $INTER \rightarrow INTRA_0$: 7; $INTER \rightarrow INTRA_1$: 5; INTRA0->INTRA1: 4 |
| Richoux FC | RICHOUX-REGULAR | 10 | 11 | |
| Richoux FC | RICHOUX-STRICT | 1 | 1 | |
| Richoux LSTM | D-SCRIPT UNBAL. | 1 | 1 | $INTER \rightarrow INTRA_0$: 1; $INTER \rightarrow INTRA_1$: 1; INTRA0->INTRA1: 1 |
| Richoux LSTM | DU | 1 | 1 | $INTER \rightarrow INTRA_0$: 1; $INTER \rightarrow INTRA_1$: 1; INTRA0->INTRA1: 1 |
| Richoux LSTM | GUO | 1 | 1 | $INTER \rightarrow INTRA_0$: 1; $INTER \rightarrow INTRA_1$: 1; INTRA0->INTRA1: 1 |
| Richoux LSTM | HUANG | 1 | 1 | $INTER \rightarrow INTRA_0$: 1; $INTER \rightarrow INTRA_1$: 1; INTRA0->INTRA1: 1 |
| Richoux LSTM | PAN | 1 | 24 | $INTER \rightarrow INTRA_0$: 1; $INTER \rightarrow INTRA_1$: 1; INTRA0->INTRA1: 1 |
| Richoux LSTM | RICHOUX | | | $INTER \rightarrow INTRA_0$: 1; $INTER \rightarrow INTRA_1$: 8; INTRA0->INTRA1: 13 |
| Richoux LSTM | RICHOUX-REGULAR | 17 | 16 | |
| Richoux LSTM | RICHOUX-STRICT | 1 | 19 | |
| D-SCRIPT | D-SCRIPT UNBAL. | 7 | 2 | $INTER \rightarrow INTRA_0$: 10; $INTER \rightarrow INTRA_1$: 4; INTRA0->INTRA1: 1 |
| D-SCRIPT | DU | 9 | 9 | $INTER \rightarrow INTRA_0$: 10; $INTER \rightarrow INTRA_1$: 10; INTRA0->INTRA1: 5 |
| D-SCRIPT | GUO | 10 | 9 | $INTER \rightarrow INTRA_0$: 6; $INTER \rightarrow INTRA_1$: 3; INTRA0->INTRA1: 2 |
| D-SCRIPT | HUANG | 5 | 8 | $INTER \rightarrow INTRA_0$: 8; $INTER \rightarrow INTRA_1$: 9; INTRA0->INTRA1: 8 |
| D-SCRIPT | PAN | 9 | 6 | $INTER \rightarrow INTRA_0$: 8; $INTER \rightarrow INTRA_1$: 2; INTRA0->INTRA1: 1 |
| D-SCRIPT | RICHOUX | | | $INTER \rightarrow INTRA_0$: 6; $INTER \rightarrow INTRA_1$: 1; INTRA0->INTRA1: 3 |
| D-SCRIPT | RICHOUX-REGULAR | 1 | 1 | |
| D-SCRIPT | RICHOUX-STRICT | 1 | 1 | |
| Topsy-Turvy | D-SCRIPT UNBAL. | 4 | 3 | $INTER \rightarrow INTRA_0$: 5; $INTER \rightarrow INTRA_1$: 7; INTRA0->INTRA1: 1 |
| Topsy-Turvy | DU | 3 | 10 | $INTER \rightarrow INTRA_0$: 1; $INTER \rightarrow INTRA_1$: 4; INTRA0->INTRA1: 5 |

Table S7: Epochs in which early stopping was triggered.

| | | | | |
|---|---|---|---|---|
| Topsy-Turvy | GUO | 7 | 9 | *INTER* $\rightarrow$ *INTRA*$_0$: 8; *INTER* $\rightarrow$ *INTRA*$_1$: 3; INTRA0->INTRA1: 3 |
| Topsy-Turvy | HUANG | 7 | 4 | *INTER* $\rightarrow$ *INTRA*$_0$: 10; *INTER* $\rightarrow$ *INTRA*$_1$: 1; INTRA0->INTRA1: 9 |
| Topsy-Turvy | PAN | 2 | 7 | *INTER* $\rightarrow$ *INTRA*$_0$: 7; *INTER* $\rightarrow$ *INTRA*$_1$: 9; INTRA0->INTRA1: 1 |
| Topsy-Turvy | RICHOUX | | | *INTER* $\rightarrow$ *INTRA*$_0$: 9; *INTER* $\rightarrow$ *INTRA*$_1$: 2; INTRA0->INTRA1: 10 |
| Topsy-Turvy | RICHOUX-REGULAR | 1 | 1 | |
| Topsy-Turvy | RICHOUX-STRICT | 5 | 1 | |

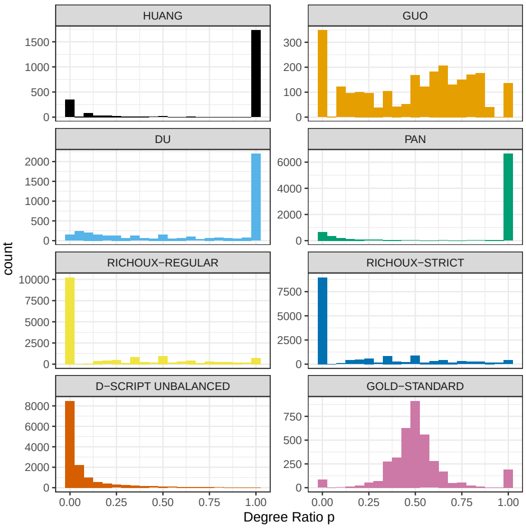


(a)

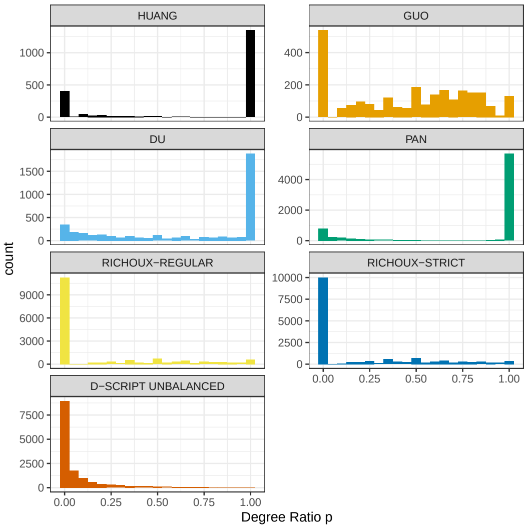


(b)

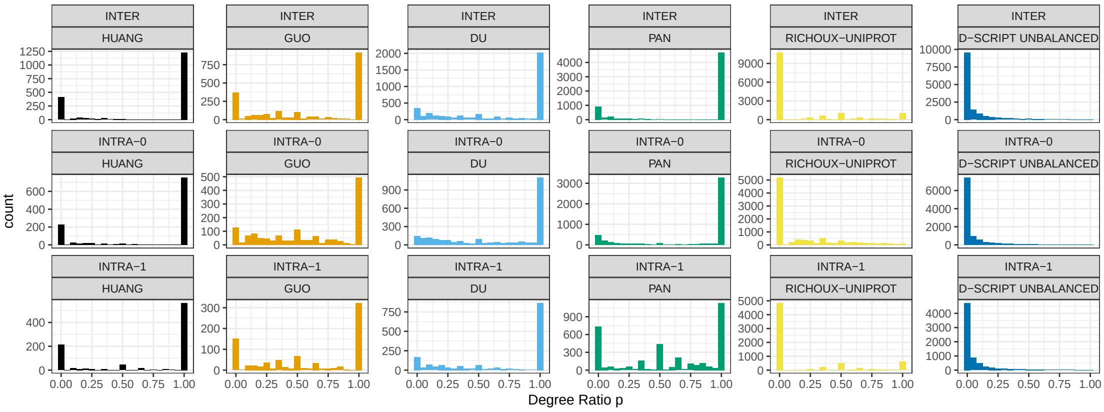


(c)

Figure S6: Degree ratios for the training datasets for the **(a)** original, **(b)** rewired, and **(c)** partition datasets. Most of the proteins in HUANG and PAN have either exclusively positive or negative interactions. DU has many proteins that only have positive interactions, the RICHOUX datasets have many proteins with only negative annotations.

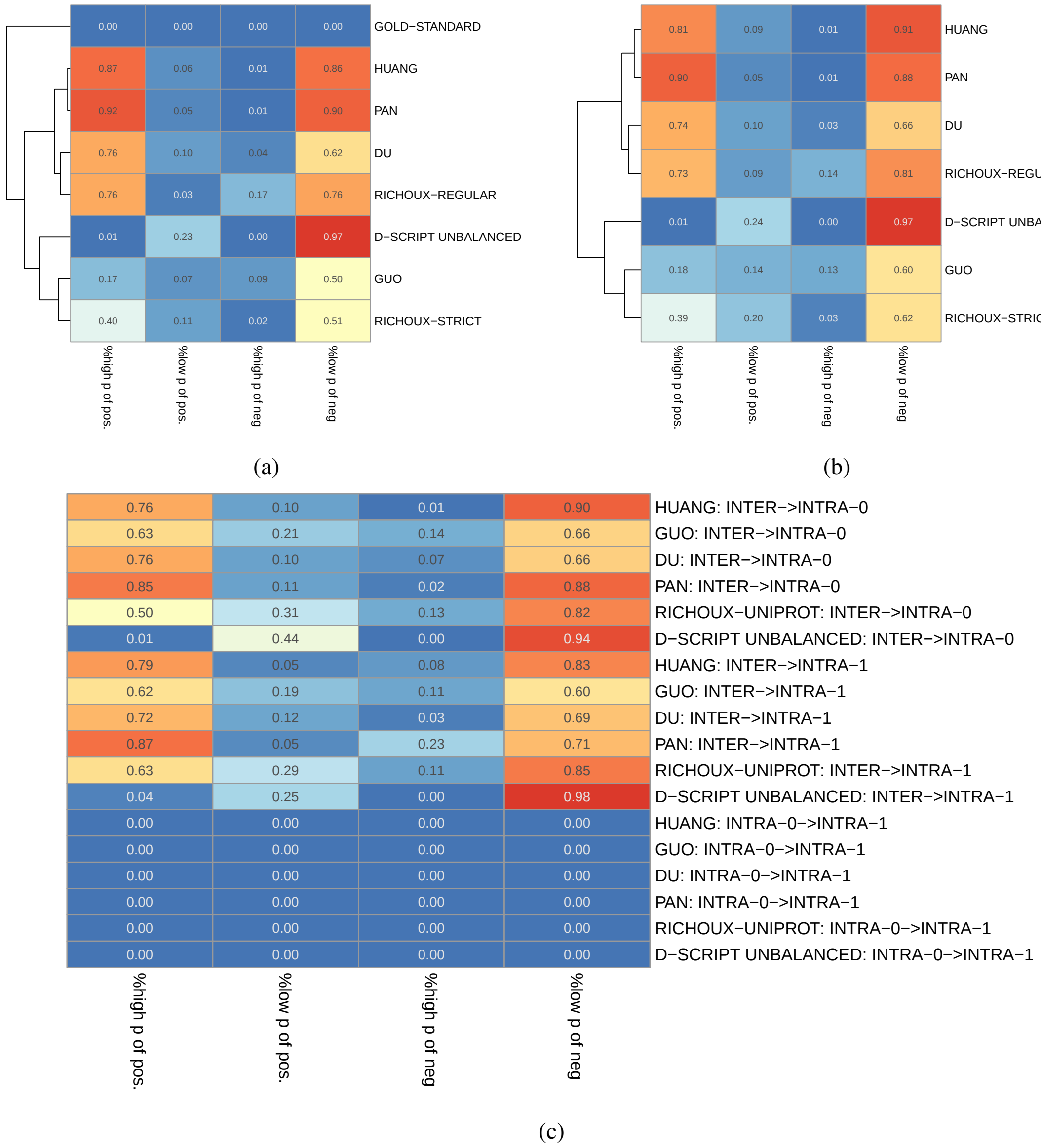


Figure S7: Proportions of training proteins with a high ($\geq 0.9$) or low ($\leq 0.1$) degree ratio in the positive and negative parts of the test sets displayed for the **(a)** original, **(b)** rewired, and **(c)** partition datasets. Consequently, especially HUANG, and PAN should be very easy to predict. RICHOUX-REGULAR and DU should be a bit harder and GUO and RICHOUX-STRICT the hardest. This trend is visible in Figure 2a and 3a

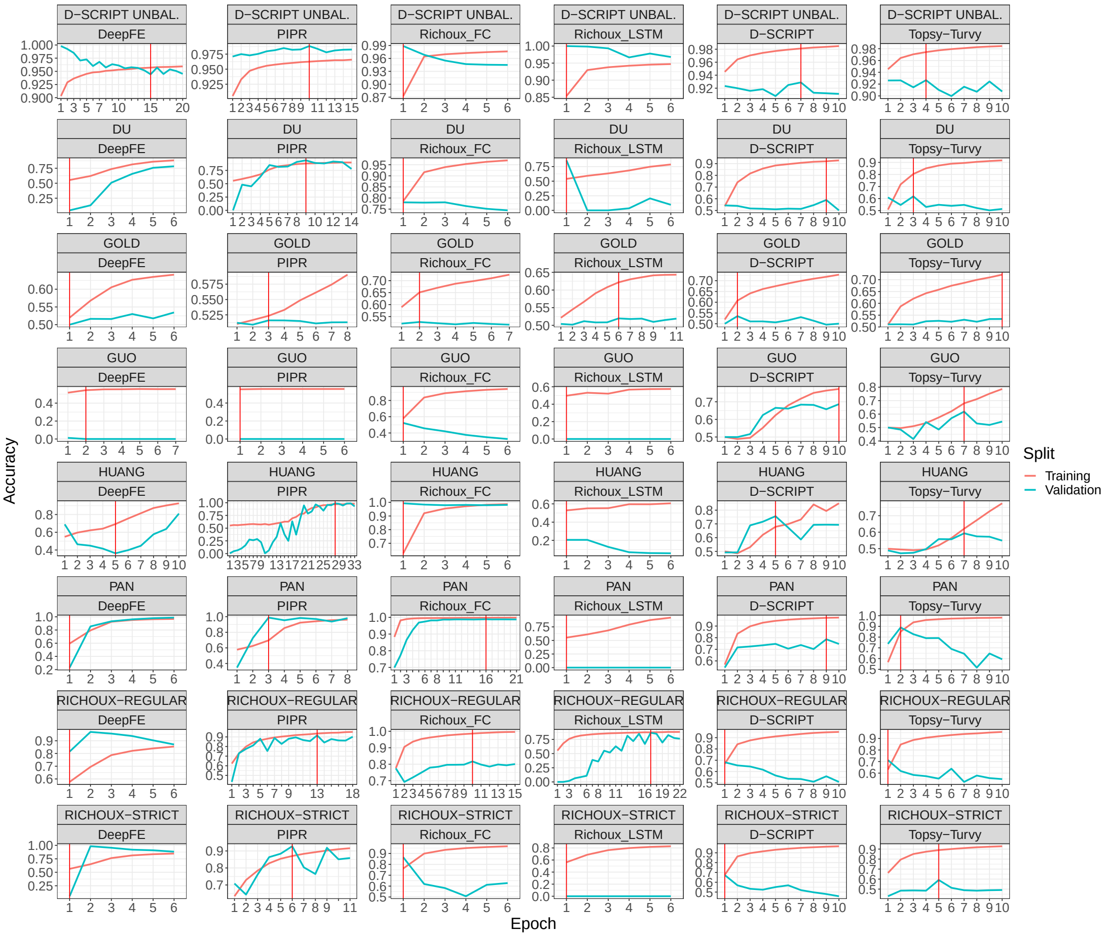


Figure S8: Training and validation accuracy over all epochs until early stopping for all deep learning methods on the original datasets. The red line indicates which model was used to evaluate the test set in the early stopping setting.

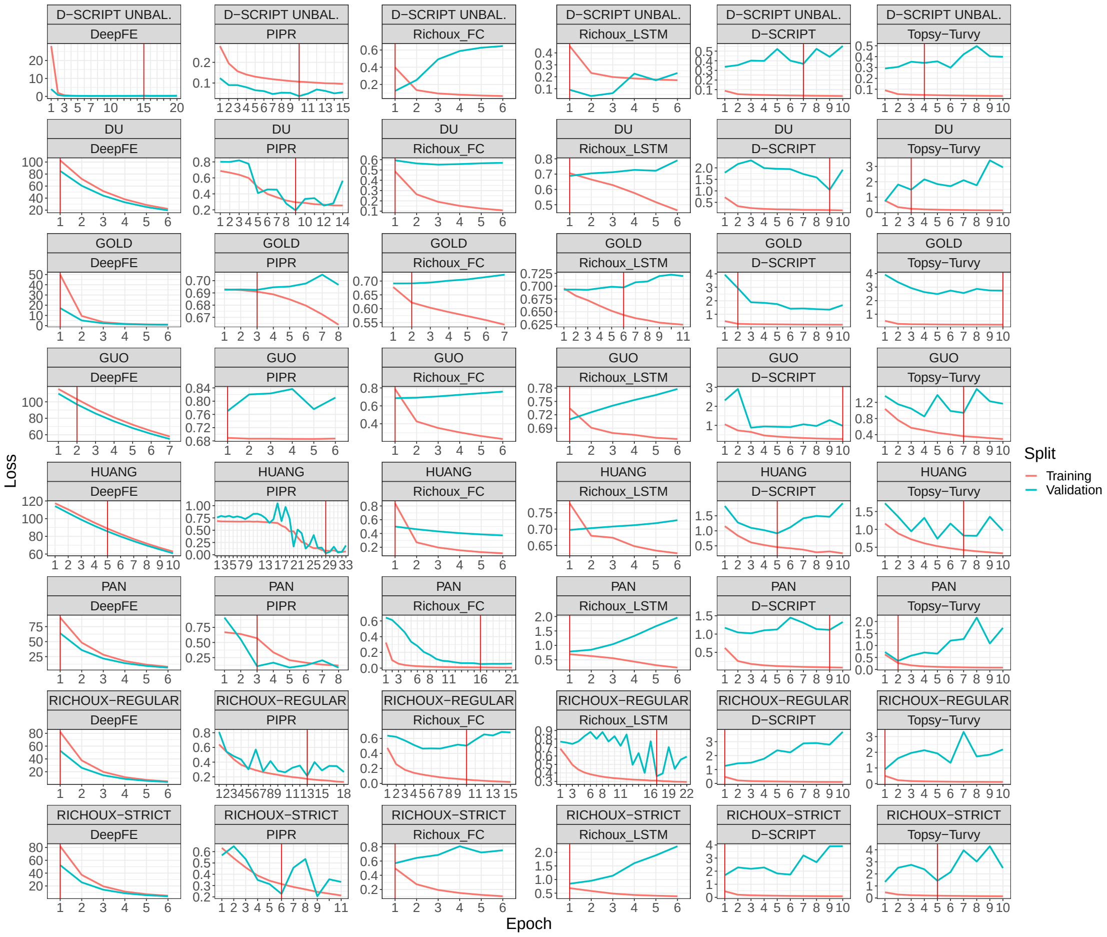

Figure S9: Training and validation loss over all epochs until early stopping for all deep learning methods on the original datasets. The red line indicates which model was used to evaluate the test set in the early stopping setting.

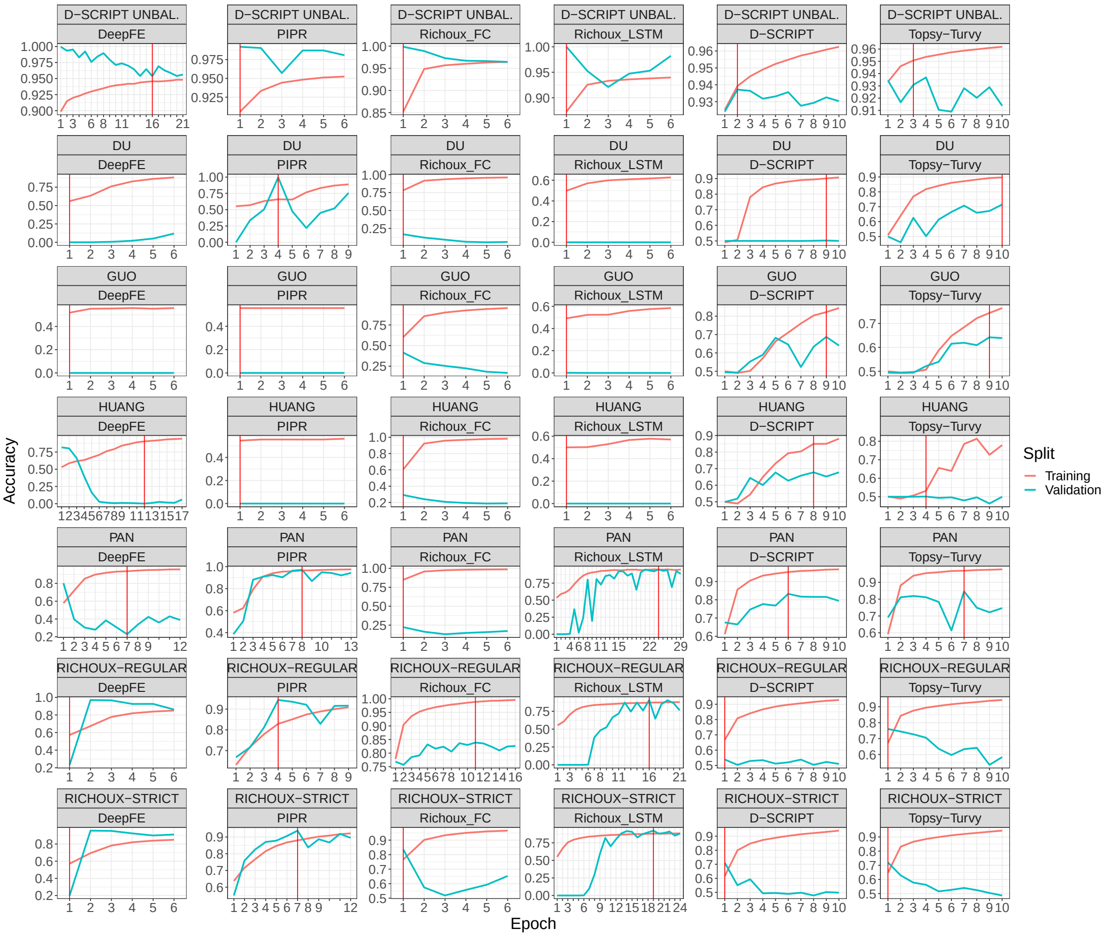


Figure S10: Training and validation accuracy over all epochs until early stopping for all deep learning methods on the rewired datasets. The red line indicates which model was used to evaluate the test set in the early stopping setting.

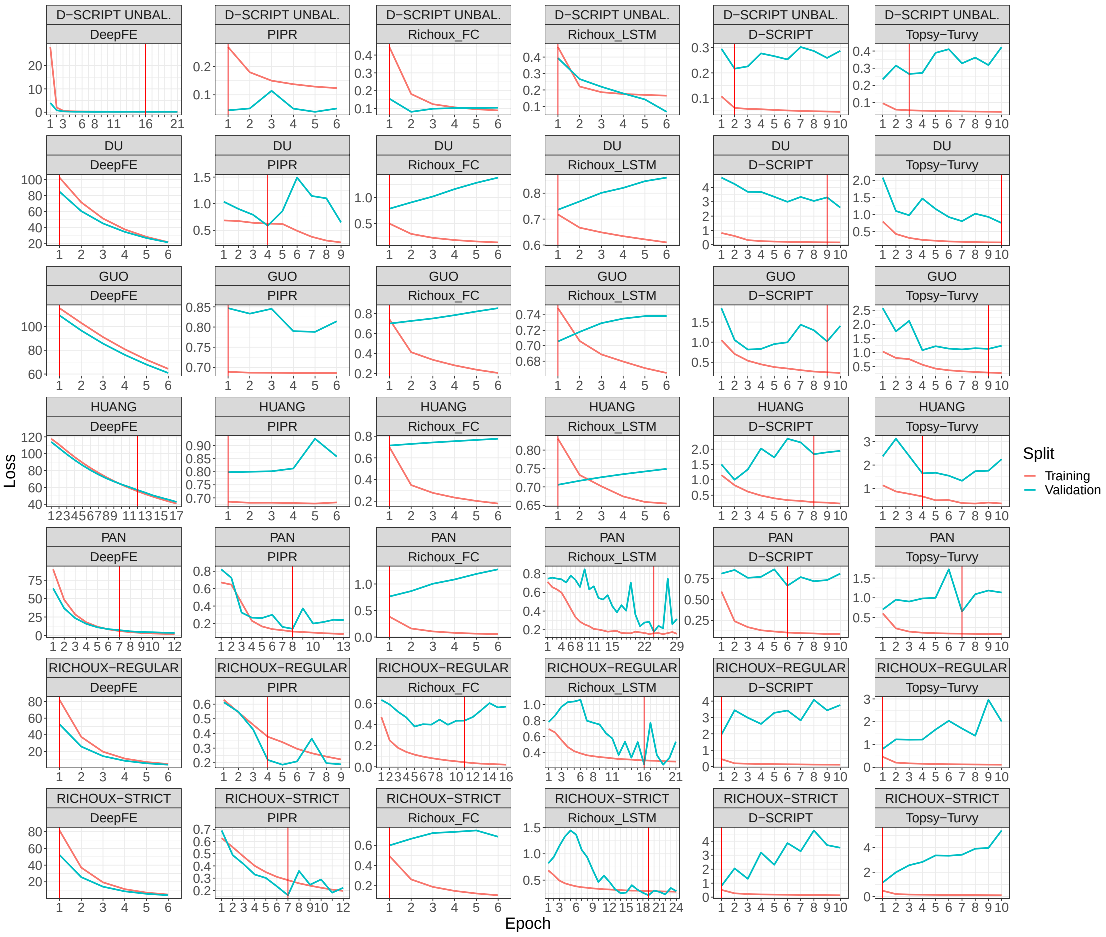


Figure S11: Training and validation loss over all epochs until early stopping for all deep learning methods on the rewired datasets. The red line indicates which model was used to evaluate the test set in the early stopping setting.

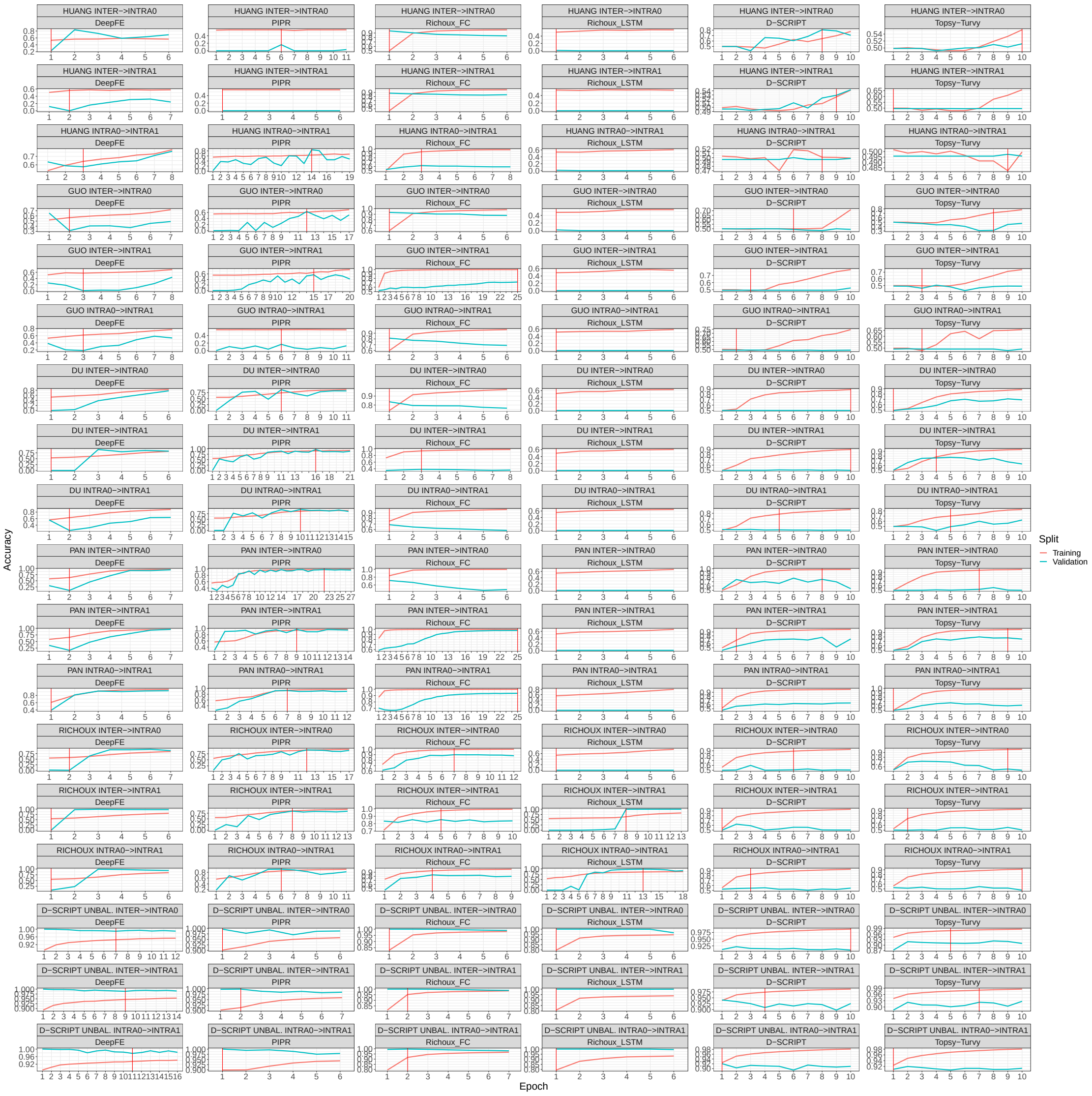

Figure S12: Training and validation accuracy over all epochs until early stopping for all deep learning methods on the partition datasets. The red line indicates which model was used to evaluate the test set in the early stopping setting.

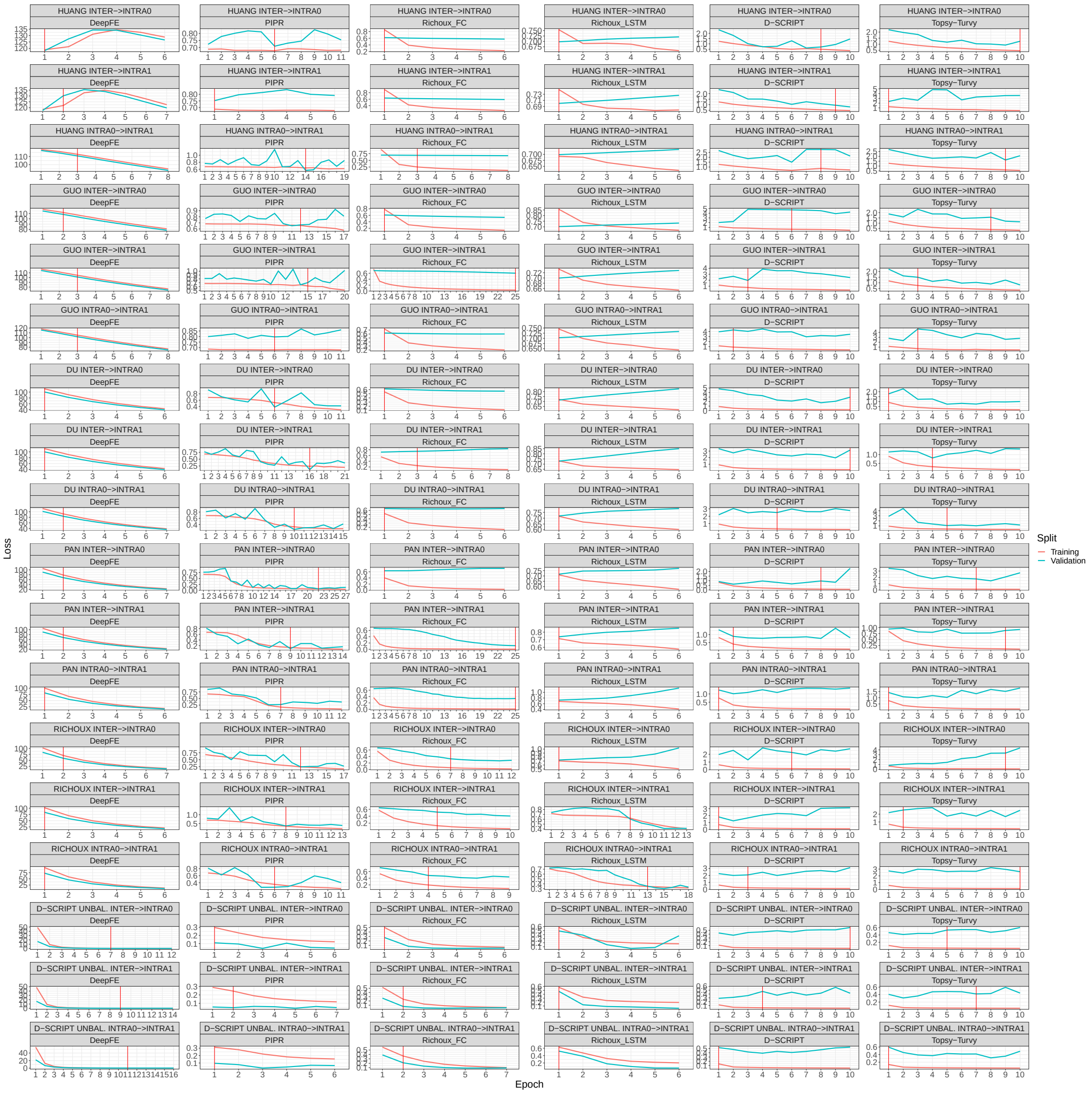

Figure S13: Training and validation loss over all epochs until early stopping for all deep learning methods on the partition datasets. The red line indicates which model was used to evaluate the test set in the early stopping setting.

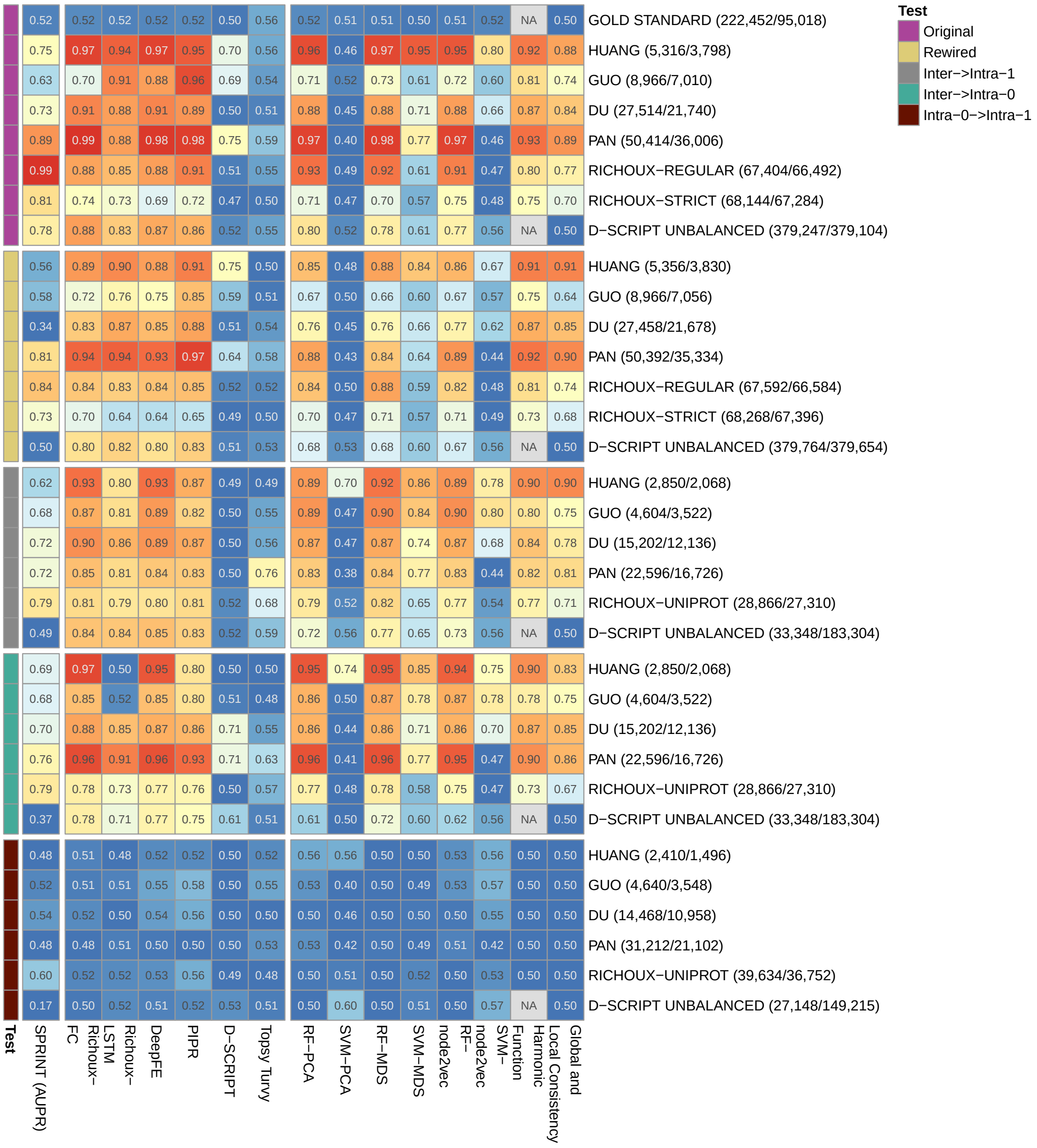


Figure S14: Balanced accuracies (AUPRs for SPRINT) for all methods and datasets.

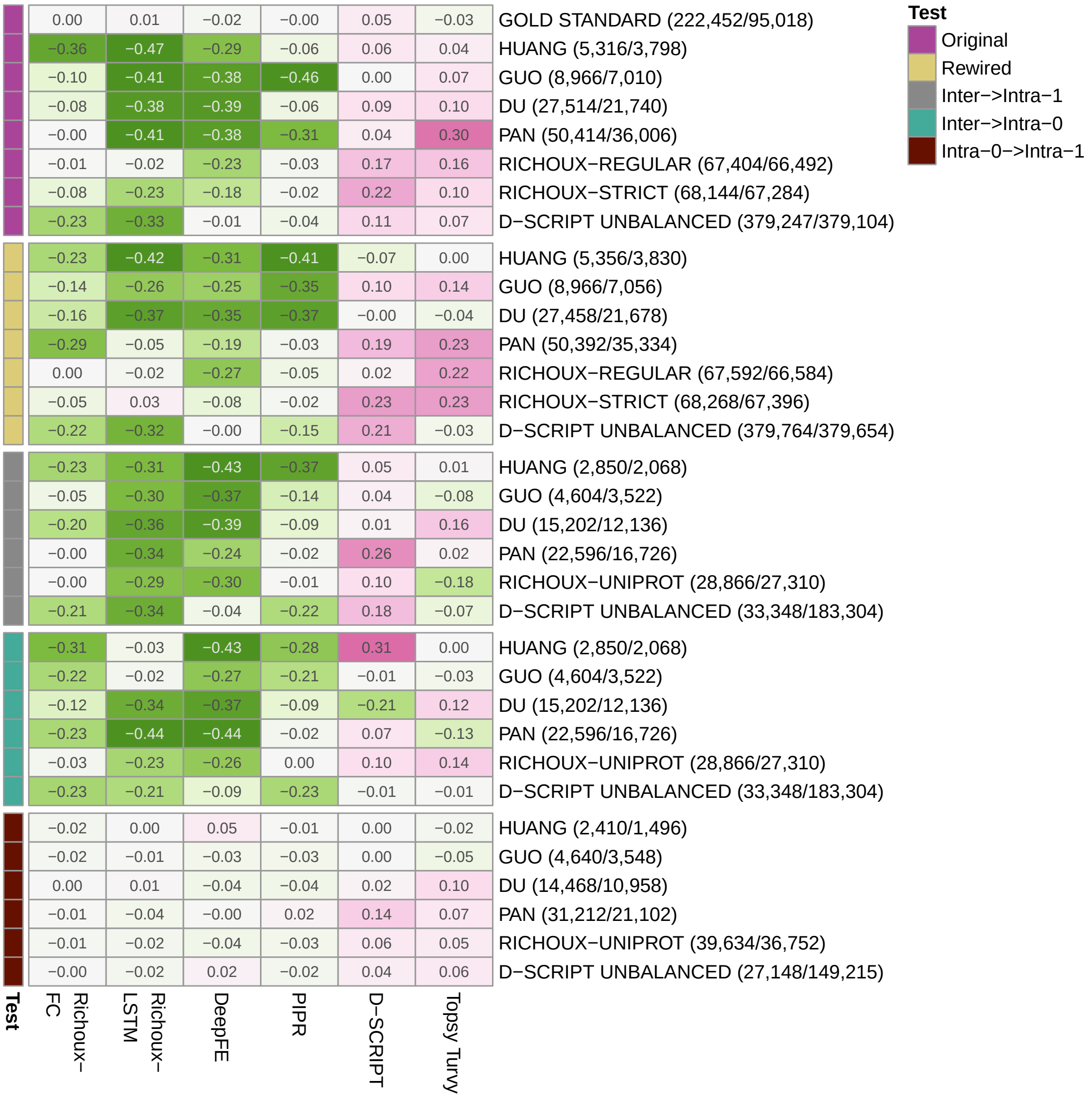


Figure S15: Difference between the balanced accuracies from the early stopping setting and the balanced accuracies obtained without early stopping. Only D-SCRIPT and Topsy-Turvy seem to profit from early stopping.

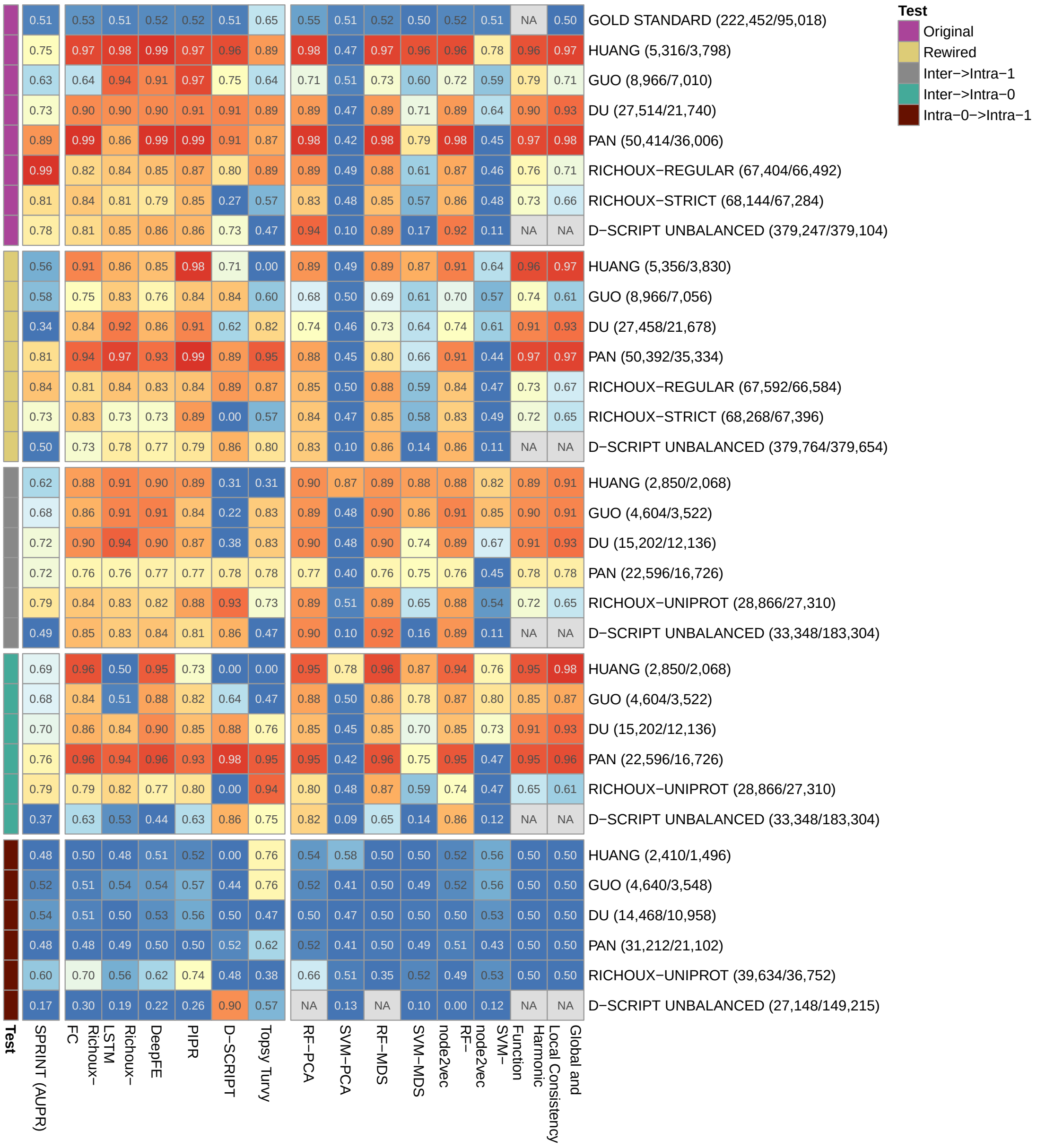


Figure S16: Precision (AUPRs for SPRINT) for all methods and datasets.

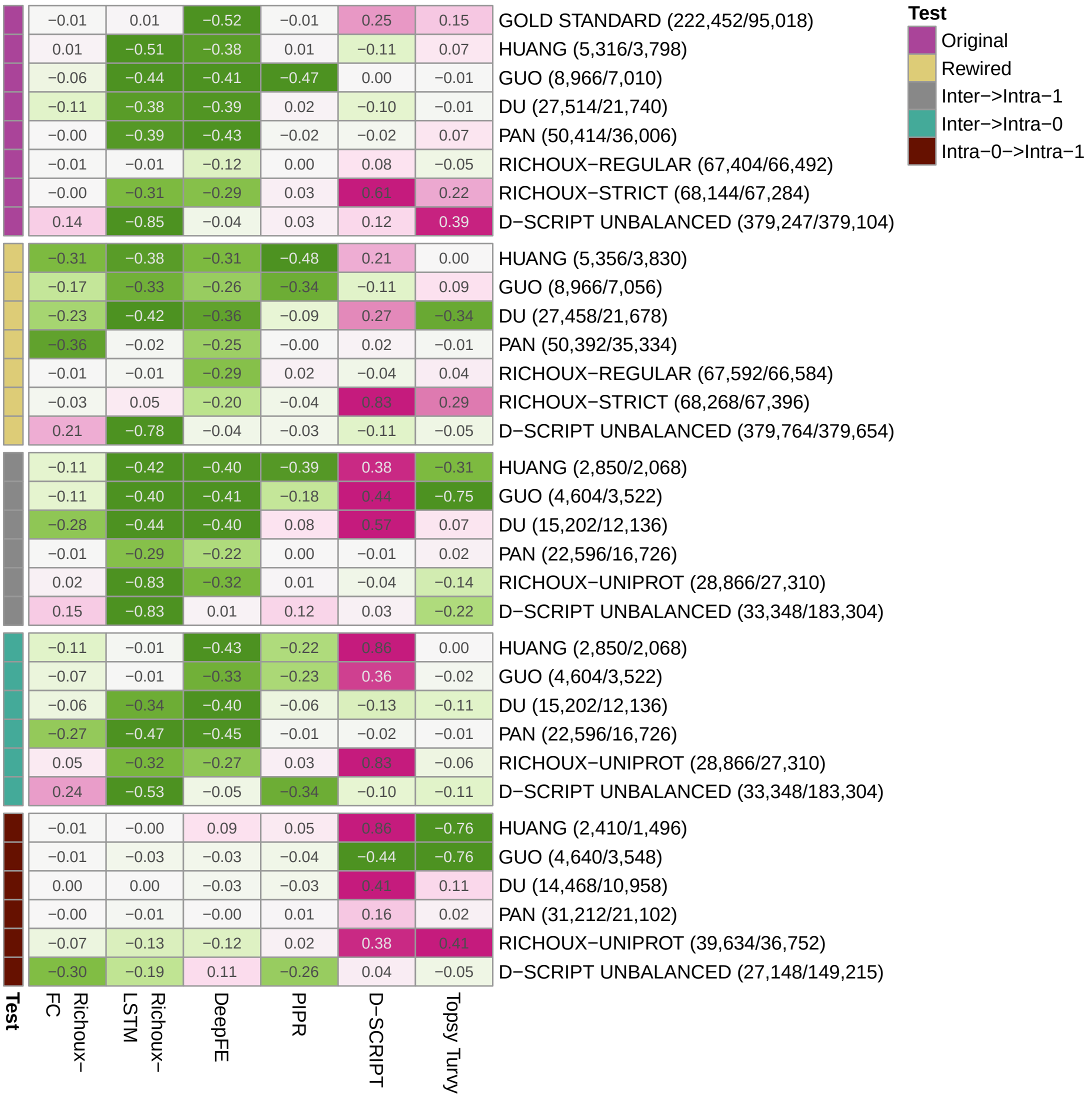


Figure S17: Difference between the precisions from the early stopping setting and the precisions obtained without early stopping. Precision increases especially for D-SCRIPT.

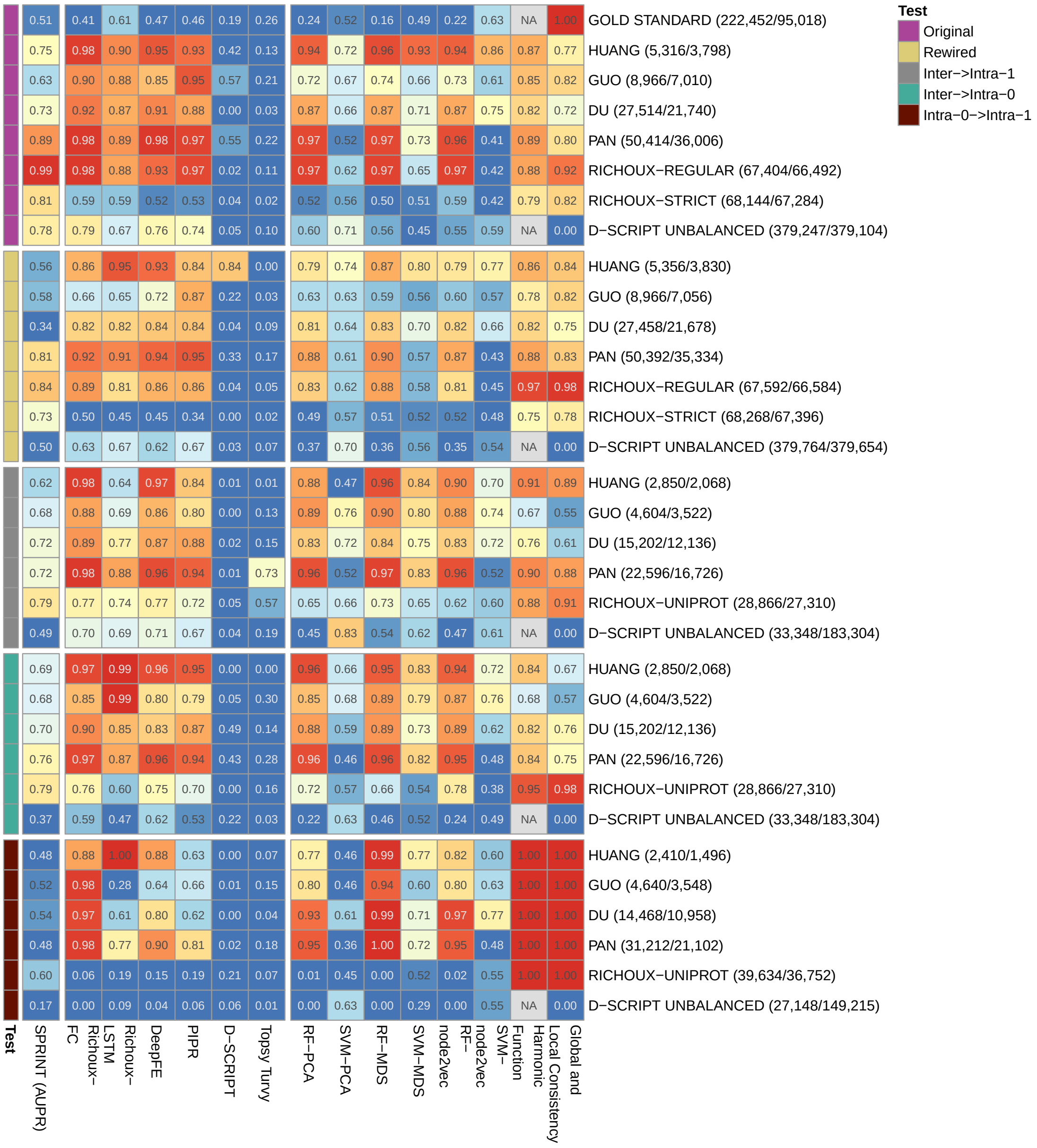


Figure S18: Recall (AUPRs for SPRINT) for all methods and datasets.

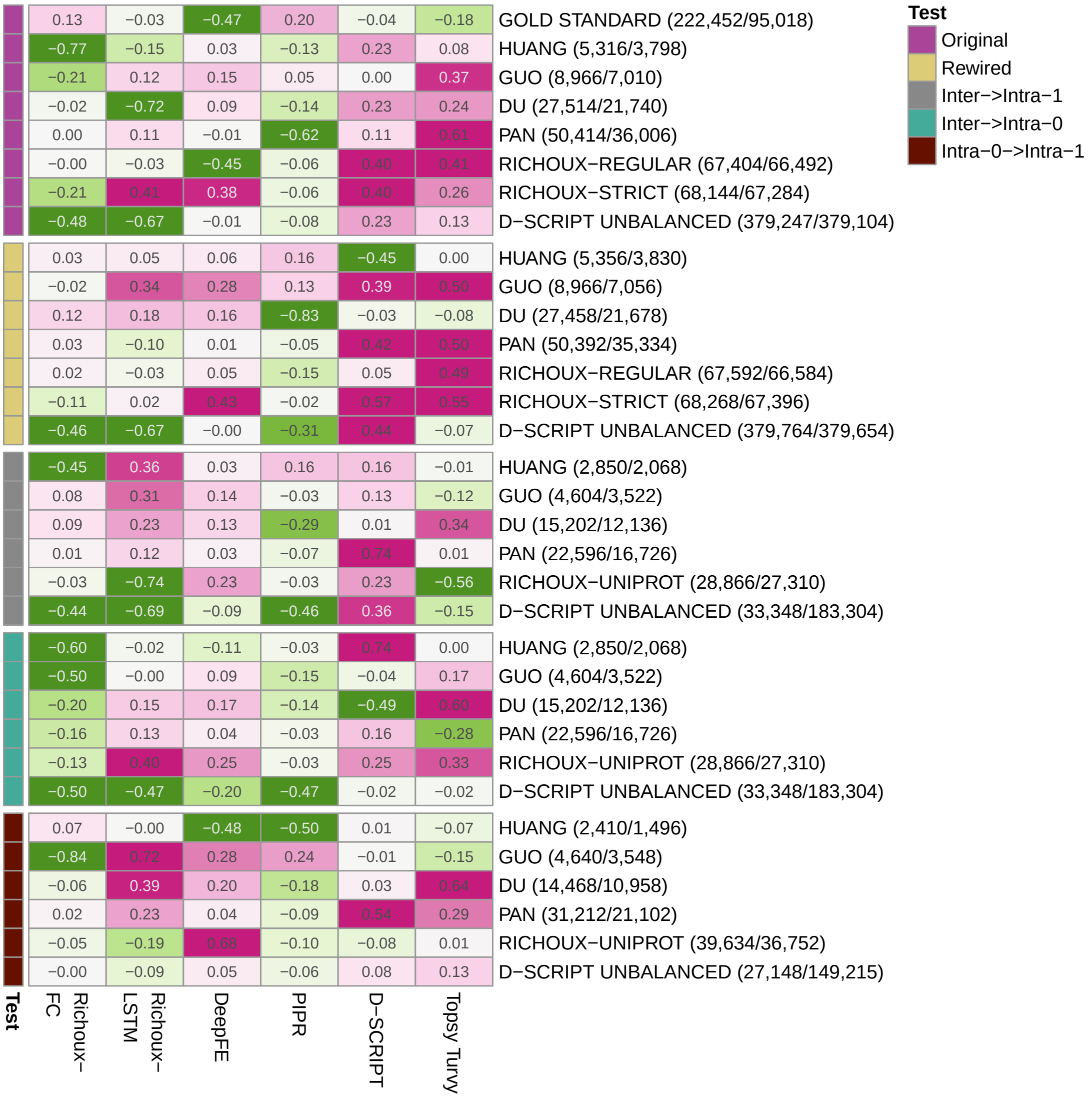


Figure S19: Difference between the recalls from the early stopping setting and the recalls obtained without early stopping. Recall increases for most methods except for Richoux-FC and PIPR.

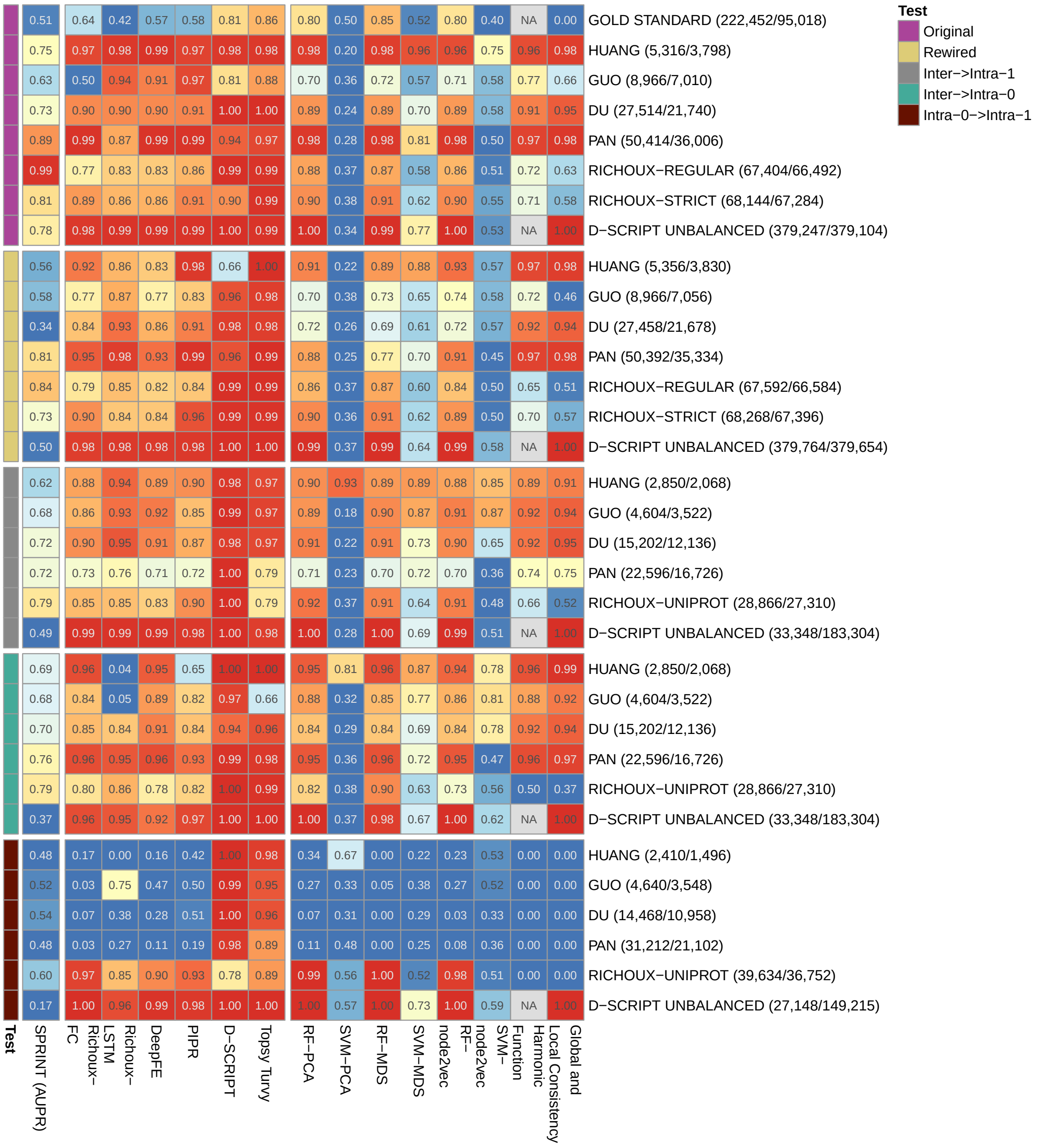


Figure S20: Specificity (AUPRs for SPRINT) for all methods and datasets.

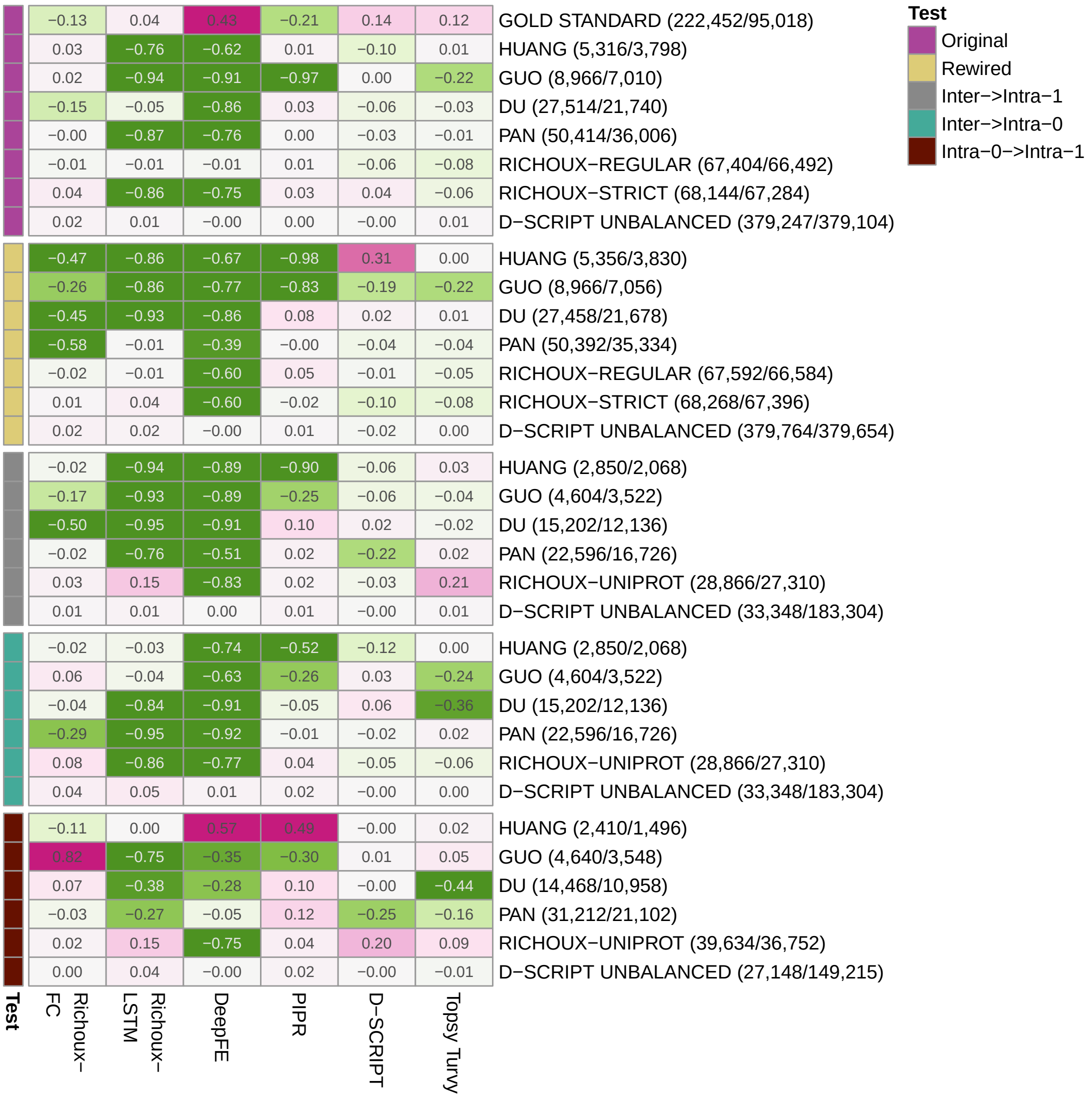


Figure S21: Difference between the specificities from the early stopping setting and the specificities obtained without early stopping. Specificity either mostly decreases (Richoux-LSTM, DeepFE) or stays approximately the same.

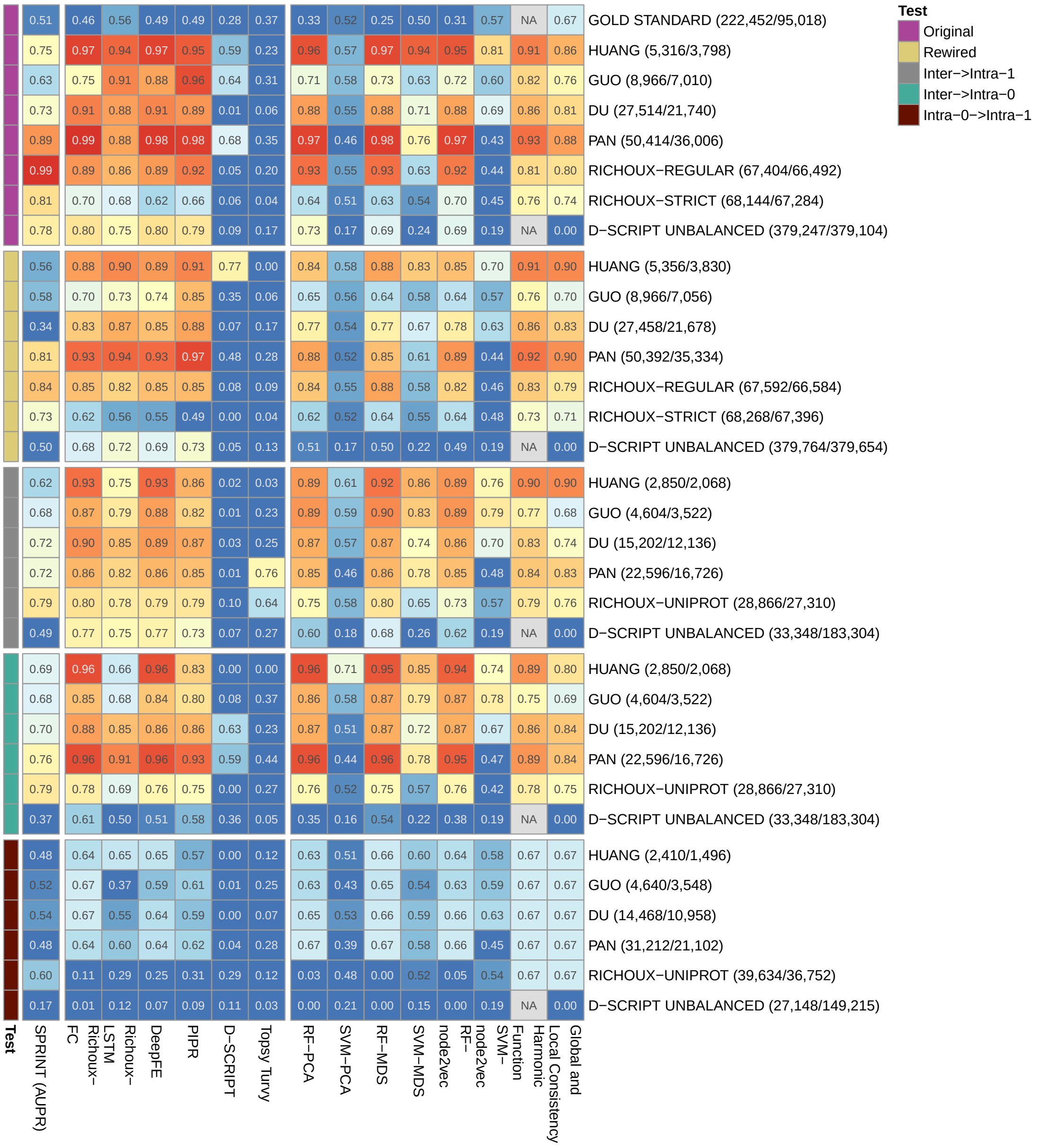


Figure S22: F1 (AUPRs for SPRINT) for all methods and datasets.

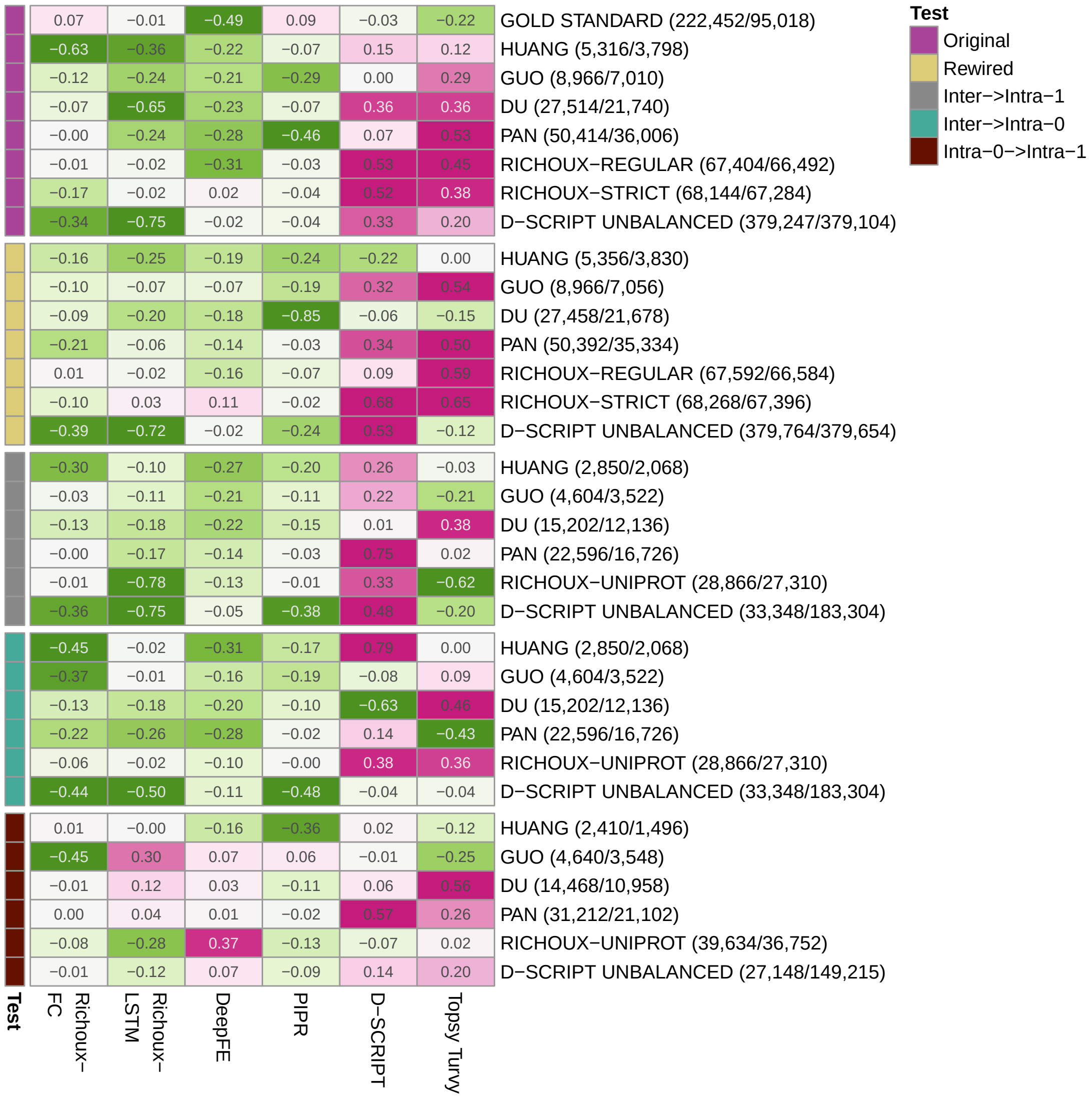


Figure S23: Difference between the F1 scores from the early stopping setting and the F1 scores obtained without early stopping. Mostly D-SCRIPT and Topsy-Turvy profit from early stopping. The performance from Richoux-LSTM and DeepFE improves for the $INTRA_0 \rightarrow INTRA_1$ setting.

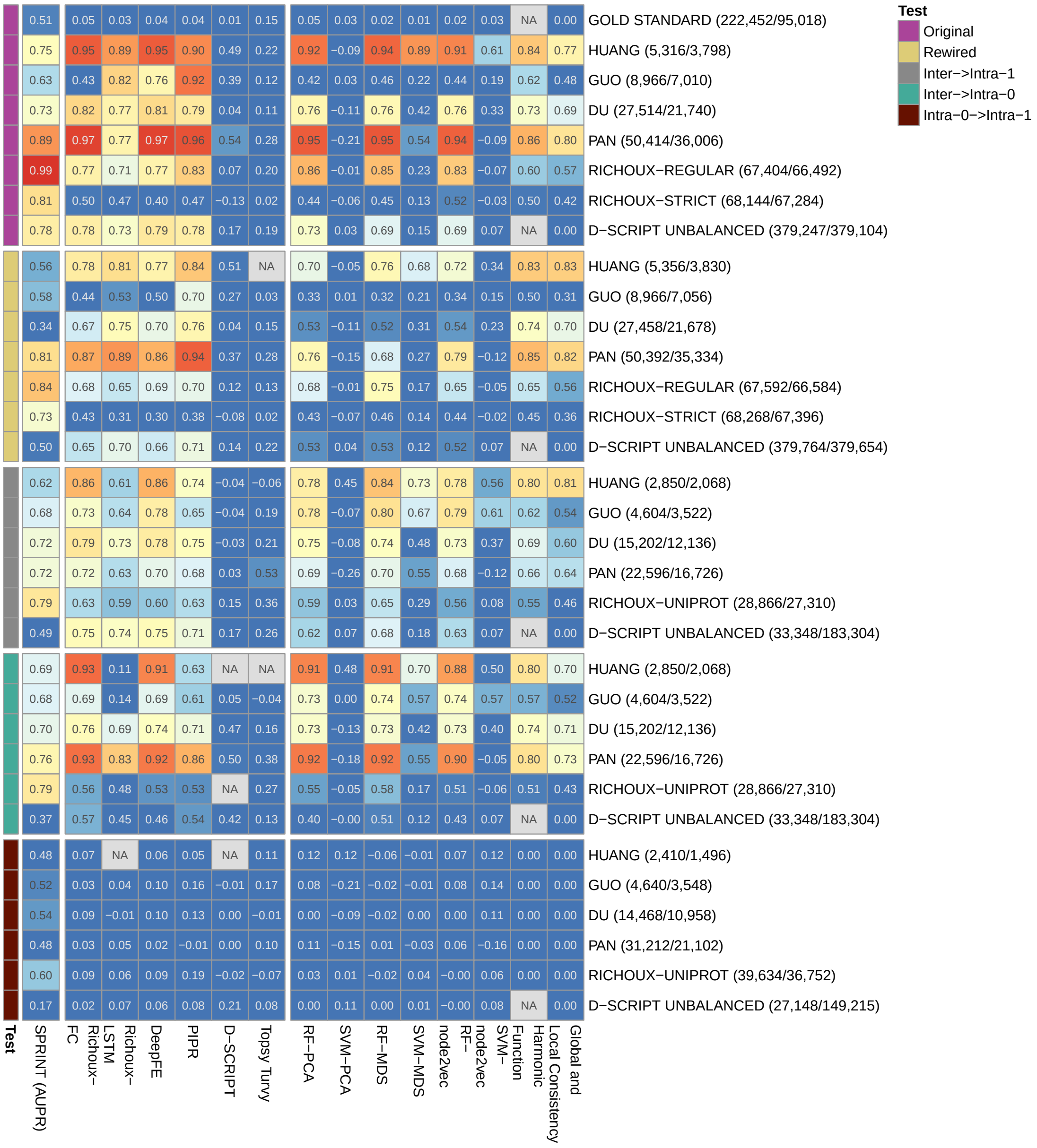


Figure S24: MCC (AUPRs for SPRINT) for all methods and datasets.

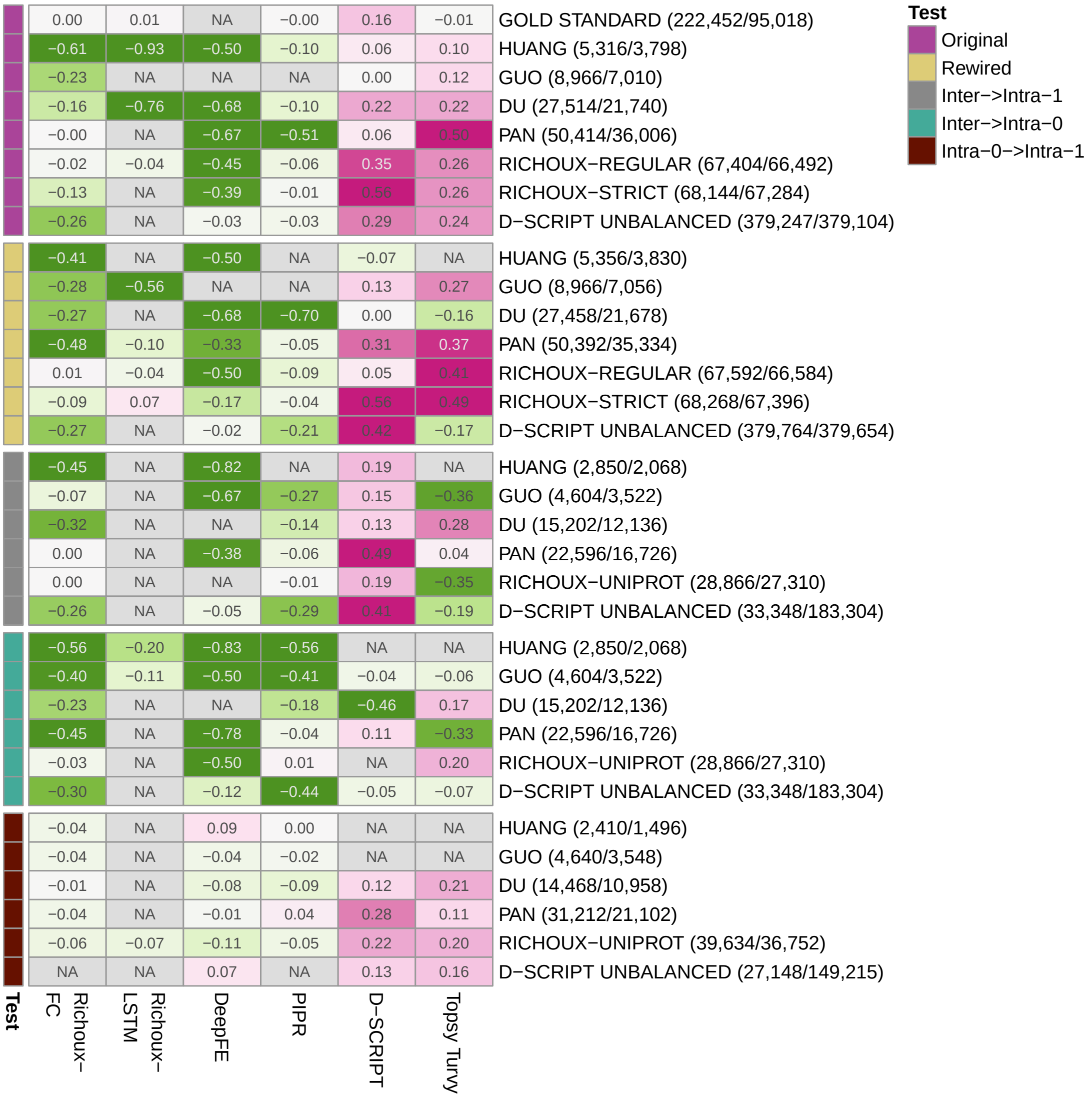


Figure S25: Difference between the MCC scores from the early stopping setting and the MCC scores obtained without early stopping. Mostly D-SCRIPT and Topsy-Turvy profit from early stopping.
